# Combining brain-wide activity imaging with electron microscopy reveals a distributed brain network for processing threatening somatosensory stimuli

**DOI:** 10.1101/2025.09.25.678485

**Authors:** Nadine Randel, Chen Wang, Michael S Clayton, Kun Wang, Song Pang, C. Shan Xu, Andrew S Champion, Harald F Hess, Albert Cardona, Philipp J Keller, Marta Zlatic

## Abstract

To understand how brains work, it is necessary to connect neural activity to synaptic-resolution circuit architecture. Recent advances in light-sheet microscopy (LSM) enable whole-brain, cellular-resolution imaging of activity of all neuronal cell bodies, however, most neurons from such datasets cannot be identified. In most organisms, neurons are identifiable based on their projections (and not based on their cell body position) which, when densely labelled, cannot be resolved using LSM. Here, we present a novel methodology to overcome this by combining whole-brain activity imaging with subsequent volume electron microscopy imaging of the same brain to visualise neuronal projections and identify neurons with interesting activity. We used this approach to identify brain neurons that process input from multisensory (nociceptive and mechanosensory) Basin interneurons that trigger vigorous escape in *Drosophila* larvae in response to threatening somatosensory stimuli. After whole-brain imaging of neuronal activity during Basin activation, we imaged the same brain with an enhanced focused ion-beam electron microscope (eFIB-SEM). We registered the functional and anatomical volumes and reconstructed (in the eFIB-SEM volume) the projections of neurons that responded to Basin activation to determine their developmental lineage and identity. This revealed a distributed network for processing threatening somatosensory stimuli that trigger vigorous escape spanning 25 distinct lineages and many distinct brain areas, and included direct brain targets of Basin neurons that integrate somatosensory information with other modalities, as well as brain output neurons (descending neurons [DN]) that likely contribute to action-selection. Our workflow provides a powerful framework for mapping neuronal activity onto structure across an entire brain, yielding novel insights into the distributed central processing of noxious stimuli.

## INTRODUCTION

In order to behave adaptively in the environment, animals must process incoming sensory information and select appropriate actions, but we still lack a comprehensive understanding of neural circuits that underlie these fundamental brain functions. One major challenge is that even seemingly simple behaviours, such as escape responses to threatening somatosensory stimuli, are subserved by circuits that are not restricted to small, specific brain regions but distributed across many areas of the central nervous system.

Our understanding of neuronal mechanisms has been greatly advanced by access to whole-brain connectomes, which reveal the patterns of connections between all neurons in the brain ^1–9^. However, while synaptic-resolution connectivity maps are essential for understanding how neural circuits implement fundamental brain functions, by themselves they are not sufficient. To gain a mechanistic understanding of circuit function, in addition to knowing their connectivity patterns, it is also necessary to characterise neuronal activity patterns during a range of behavioural tasks. Indeed, numerous previous studies have shown the importance of linking both synaptic connectivity and neuronal activity on a single cell level to gain a detailed mechanistic understanding of neural circuit mechanisms ^10–24^.

However, most studies use genetic tools to selectively label and analyse response properties of one identified neuron type at a time. By combining recordings from different neurons, populations, or regions from many different animals during the same task, information about response properties from different neurons can slowly be accumulated. The problem with this approach is that it is very slow, and it would take a very long time to obtain information from all neuron types, during the same task. Furthermore, correlations in activity between different neuron types in the same individual cannot be properly assessed.

Recently, advances in light-sheet microscopy (LSM) have opened doors to imaging activity of all neurons in the entire brain of relatively small animals with single cell resolution, including nematodes, insects, and even small vertebrates ^25,26^. Such datasets can comprehensively reveal all neuronal cell bodies with specific response properties. However, in most brains larger than *C. elegans* (where such studies have been extremely fruitful), including those of small insects, like *Drosophila* larvae, cell body position is insufficient to identify neurons ^27–29^. It is, therefore, impossible to combine information about activity of specific neurons from such datasets with previously available information about these neurons–such as their developmental origin (lineage), projection pattern, synaptic connectivity, or molecular properties. In contrast to cell body position, most neurons can be uniquely identified based on the shape and location of their projections. However, while LSM provides sufficient resolution to resolve densely labelled cell bodies from each other (each about 5 µm in diameter), it does not provide sufficient resolution to resolve densely labelled neuronal projections in neuropil areas.

We have therefore developed a methodology that enables us to visualise the projections of all neurons after imaging their activity with LSM, in order to identify neurons based on their projections and combine this information with known information about their connectivity and other properties. Our approach involves imaging the same nervous system with electron microscopy (EM) after whole-brain imaging of neural activity. EM can provide sufficient resolution to resolve densely labelled projections of all neurons in the nervous system, and identify neurons based on their projection patterns after segmenting large axonal and dendritic branches. Furthermore, recent advances in EM enable imaging entire brains of small animals relatively rapidly. For example, a brain of a *Drosophila* larva (100 µm x 100 µm x 80 µm) can be imaged in one go, in under ten days, with 8 x 8 x 8 nm/voxel resolution with a single enhanced focused ion-beam electron microscope (eFIB-SEM), and other methods allow for even faster imaging of larger brains ^30–32^.

We applied this approach to identifying novel elements of brain circuits that process threatening somatosensory stimuli in the tractable insect model system, the *Drosophila* larva. The *Drosophila* larva is an excellent model system for brainwide analysis of neural circuit structure and function and for elucidating the circuit mechanisms of fundamental brain functions. Larvae have a rich behavioural repertoire, the complete synaptic-resolution connectome of a *Drosophila* larval brain has recently been published, and a large genetic toolkit is available for selectively targeting and manipulating individual neuron types and elucidating their roles in behaviour ^13,15,16,33–39^. Larval somatosensory circuits have been extensively studied and multiple neurons that respond to noxious and mechanosensory stimuli and mediate escape responses to threatening somatosensory stimuli have been identified in the ventral nerve cord ^13,40–43^. Four multisensory (nociceptive and mechanosensory) interneurons per hemisegment (Basins1-4) were shown to mediate the most vigorous escape (rolling) ^13^. Only the most threatening noxious stimuli (e.g. predator attack or contact with a very hot object) trigger the energetically costly rolling behaviour. In nature, such noxious stimuli occur in combination with mechanical stimuli. Consistent with this, combined activation of nociceptive sensory and mechanosensory neurons, induces more rolling than activation of nociceptive sensory neurons alone ^13^. Individual Basins synergistically integrate direct synaptic input from nociceptive sensory and mechanosensory and information from different Basins is synergistically integrated again by downstream neurons, so that activating all Basins 1-4 together induces significantly more rolling, than either Basin subtype alone ^13^. Thus, activating Basins 1-4 triggers more rolling and induces stronger associative learning than activation of nociceptive neurons alone ^15^. The information from Basins 1-4 is relayed to the nerve cord command neurons that trigger rolling (Gorogoro) and to the brain via ascending projection neurons that facilitate rolling ^9,13^. However, the number and types of neurons in the brain that respond to Basin 1-4 activation, their lineage identity, and whether and how they contribute to rolling is unknown.

Here, we first imaged calcium transients of all neurons in the larval brain using multi-view LSM while stimulating Basins1-4 in one nerve cord segment (A4). Subsequently, we imaged the same brain with an eFIB-SEM (imaged at 12 x 12 x 12 nm/voxel resolution), registered the LSM and eFIB-SEM volumes of the same brain to each other, and reconstructed the large projections of all neurons that had significant responses to noxious cues, to reveal their developmental origin (lineage identity), and then traced finer branches for a subset of them to uniquely identify them. We then validated our approach by using GAL4-lines to selectively express GCaMP in the identified brain neurons and confirm they respond to noxious stimuli, as revealed by pan-neuronal imaging of activity ^37^.

Our unbiased comprehensive approach provided novel insights into the brain somatosensory circuits downstream of Basins. We found about 4% of brain neurons responded significantly to activation of Basins, most of which were previously not known to respond to somatosensory stimuli. The newly identified neurons that responded to Basin stimuli spanned a range of neuron categories, including brain neurons that receive input from somatosensory projection neurons, brain output neurons (descending neurons [DN] that project to the suboesophageal zone [SEZ] and the ventral nerve cord [VNC]), as well as neurons presynaptic to the brain output neurons (pre-DN). Surprisingly, we also found that some neurons previously implicated in processing olfactory stimuli (lateral horn neurons, [LHNs]) and some neurons of the higher-order learning circuit implicated in processing conditioned stimuli (a subset of mushroom body Kenyon cells [KCs] and mushroom body output neurons [MBONs]), responded to Basin activation.

In summary, this study reveals novel insights into the distributed processing of threatening somatosensory stimuli that induce vigorous escape in the insect brain. The lineages and neurons identified here provide a valuable resource for future studies of the way in which threatening somatosensory cues are integrated with other sensory modalities and used to guide behaviour. Furthermore, the methodology developed in this study will enable overlaying activity patterns during a range of different behavioural tasks onto the connectome and provides a basis for a comprehensive brainwide understanding of neural circuit function.

## RESULTS

### Methodology for overlaying brainwide activity and connectivity maps

In order to identify brain neurons with specific functional response properties in an unbiased and comprehensive way, we developed a methodology for overlaying brainwide activity and connectivity maps (Figures 1A-J, 2A-Y and 3A-T).

**Figure 1:**
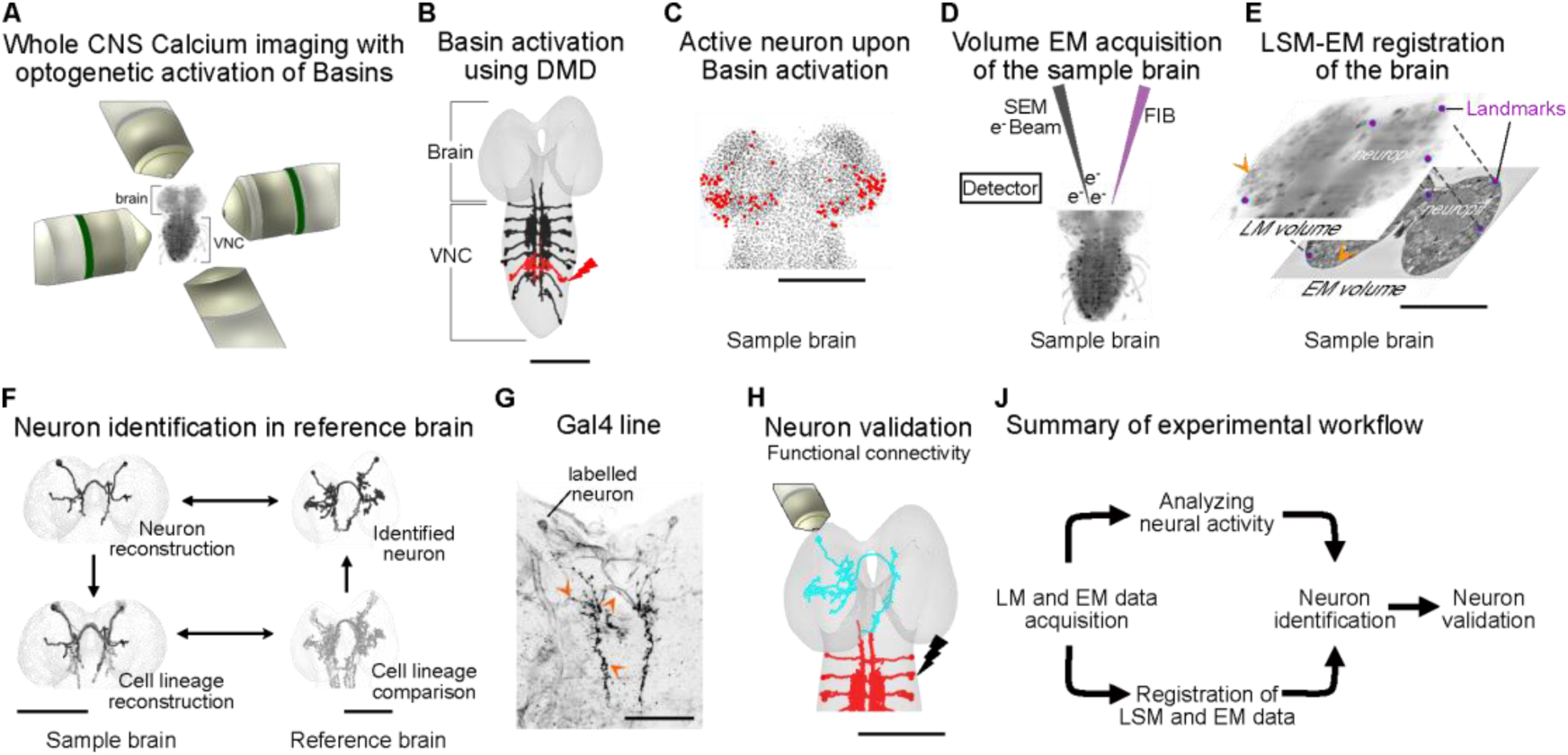
Linking single-neuron activity with synaptic connectivity. **A:** Whole central nervous system functional imaging with selective optogenetic activation, using multi-view light-sheet microscopy (SiMView). **B:** Optogenetic activation of Basins in the fourth abdominal segment using a digital micromirror device (DMD) coupled into the SiMView microscope. **C:** Location of brain neurons (red) that respond to Basin stimulation. **D:** Volume electron microscopy acquisition of the sample with known neuronal activity from panel A. **E:** Landmark-based registration of light microscopy (LM) and electron microscopy (EM) volumes of the same sample. **F:** Workflow of neuron identification in the sample brain, using the whole-brain reference connectome ^9^. **G:** Genetic driver line of the identified neuron. Arrowheads: Neurite of labelled neurons. **H:** Validation of identified neurons using 2-photon imaging of single neurons with optogenetic activation of Basins. **J:** Workflow to integrate and validate single-cell neuronal activity with synaptic connectivity. Scale bar: 50 µm.

We expressed a calcium indicator of neural activity (RGECO1a) ubiquitously in the cytoplasm of all neurons, and the optogenetic activator of neural activity (Chronos), selectively in a subset of interneurons in the nerve cord, called Basins, which receive direct synaptic input from nociceptive and mechanosensory neurons and whose optogenetic activation evokes a strong escape response ^13,44^. We imaged the activity of every neuron in the extracted nervous system (to reduce motion and deformation artefacts) of a first instar larva with a multi-view LSM, before, during and after optogenetic activation of Basins 1-4 in the fourth abdominal segment (Figure 1A-B). We presented a total of 18 Basin activations with 200 seconds between each stimulation.

Immediately after the functional experiment, we processed the sample for volume EM acquisition, by chemically fixing it, followed by staining using an adapted OTO protocol with lead aspartate to enhance contrast . We then imaged the entire CNS with eFIB-SEM at 12 x 12 x 12 nm resolution. This resolution is lower than the 8 x 8 x 8 nm resolution used to image adult fly brains, but we reasoned it could be enough to allow tracing of thicker, proximal (closer to the cell body) neuronal projections (even if not thinner, more distal branches and synapses) ^45^. After acquiring the eFIB-SEM volume, we registered the datasets of the two imaging modalities. Fixing and preparing the sample for EM imaging introduces a shrinking in size and non-linear deformations. We therefore performed a non-rigid, landmark-based deformation method for registration, the thin plate spline (TPS) transformation. We used 51 landmarks that are detectable in both volumes and evenly spaced throughout the brain to take into account non-linear deformations and ensure the method is robust (Figure 1D-E, 2A). We quantified the registration error using leave-one-out cross-validation (LOOCV, see Methods for details) and found it is highly accurate (root mean square error across each dimension is X = 1.24 µm, Y = 1.86 µm, Z = 1.86 µm). Given that the width of a soma in the *Drosophila* larva is approximately 5 µm, these sub-cellular error margins confirm that the global transformation reliably and consistently maps EM coordinates to locations in light imaging space with cellular precision.

Based on this registration, the segmented nuclei in the EM volume were projected into the LSM volume to extract their changes in fluorescence at single cell resolution (Figure 2B). Consequently, we were able to locate nuclei in the EM volume whose activity correlated with the optogenetic manipulation of Basin neurons (Figure 2C). We limited our analysis of calcium signals to cell bodies, since calcium signals from individual neurites could not be resolved in the densely labelled sample.

**Figure 2:**
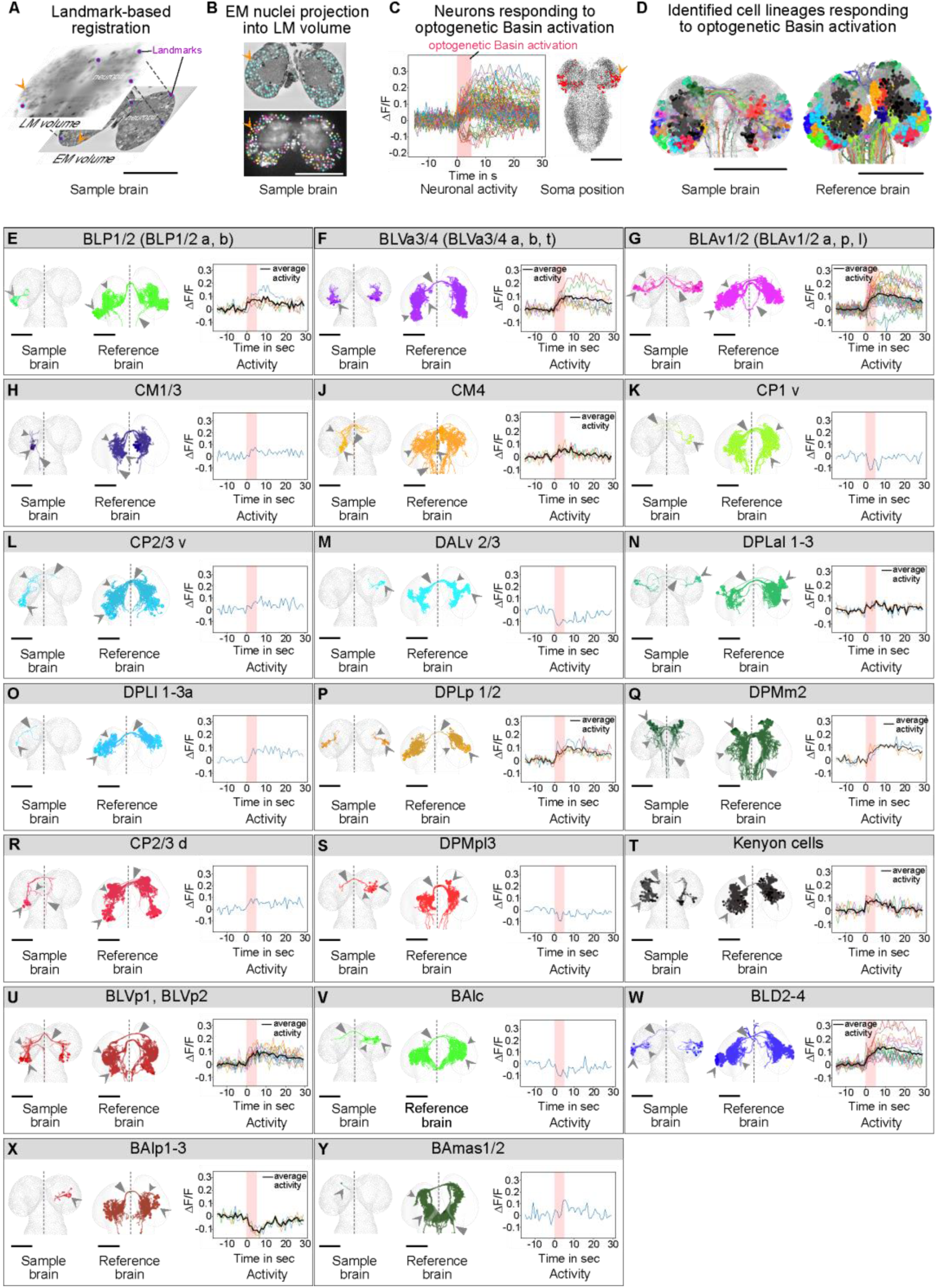
Cell-lineages responding to Basin activation. **A:** Landmark-based registration of light microscopy (LM) and electron microscopy (EM) volumes of the same sample. **B:** Automatic nuclei segmentation in the electron microscopy volume, and nuclei projection into the registered light microscopy volume. **C:** Activity traces and brain neuron locations that respond to Basin activation (red shading). **D:** Cell lineages responding to Basin stimulation in the sample brain and reference brain. Non-responding lineages are shown in grey. **E-Y:** Individual cell-lineages shown in both the sample and reference brain, with corresponding neuronal activity. Average traces shown in black. Red shading indicates time of optogenetic Basin activation. **G:** Hyperpolarised neurons are shown in a lighter shade. Dashed line indicates the midline, barbed arrowhead shows cell body, triangle indicates axon, and curved arrowhead shows dendrite. Scale bar: 25 µm. See also Figure S1 and Table S1, S4.

From the ∼3,000 brain neurons, 119 neurons (∼4%) responded consistently and significantly to the activation of Basin neurons (Figure 2C). Interestingly, the cell bodies of many of these neurons were located in the same brain region, as the cell bodies of brain neurons that receive shortest possible, two-hop input from Basins (via direct input from ascending projection neurons directly postsynaptic to Basins) in our recently-published reference brain connectome (Figure 3B) ^9^.

**Figure 3:**
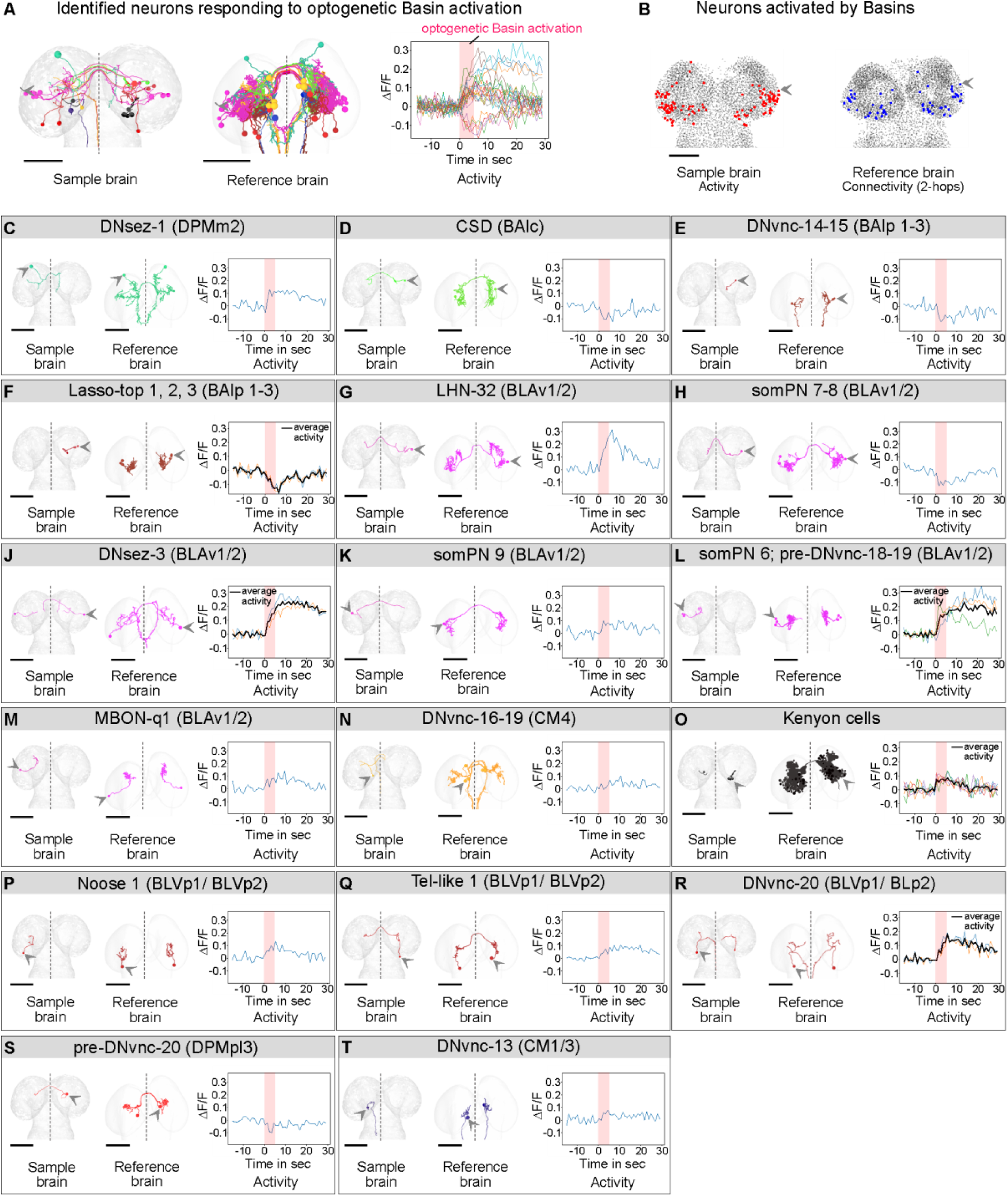
Identified neurons responding to Basin activation. **A:** Summary of identified neurons in the sample brain and reference brain, with corresponding neuronal activity. Dashed line indicates the midline. **B:** Many neurons responding to Basin activation in the sample brain (left) have similar cell body locations to brain neurons that receive shortest possible, 2-hop input from Basins in the reference connectome. **C-T:** Individual identified neurons shown in both the sample and reference brains, with corresponding neuronal activity. **E,F,H,L,N:** Candidate neurons. Individual identification was not possible due to high neuronal similarity or incomplete reconstruction. Average activity traces shown in black. Barbed arrowhead shows cell body, triangle indicates axon, and curved arrowhead shows dendrite. Scale bar: 25 µm. See also Figure S2-S3 and Table S2, S4.

To identify the developmental origin (lineage) of these neurons we manually reconstructed their thick, proximal projections up to their entry point into the neuropil and a little beyond (Figure 2D). Based on their neuropil entry point, we identified 25 neuronal lineages in which 1–30 neurons responded to Basin activation (Figure 2E-Y, Table S1).

Next, we randomly selected 50 of the 119 neurons that responded to Basin activation (Figure 3A) to determine what fraction of them could be uniquely identified based on the intermediate, more distal branches that were traceable in this 12 x 12 12 nm resolution eFIB-EM dataset. For this purpose, we traced their thinner branches as far as possible, and then compared the overall shapes of their projections to the shape of the reference neurons from our recently published, complete reconstruction of all neurons in a larval brain [and their synaptic connections] ^9^. We were able to trace intermediate-sized branches for 26/50 neurons (Figure 3A-T), whereas the intermediate-sized branches were not traceable for the remaining 24 neurons, due to the chosen 12 x 12 x 12 nm resolution of eFIB-SEM imaging. We were able to uniquely identify 18/26 individual neurons based on their intermediate-sized branches and 8/26 neurons could be narrowed down to 2 - 4 candidate neurons within the same lineage due to the high morphological similarity of their intermediate-sized branches (Figure 1F, 3C-T). Having identified neurons responsive to Basin activation, we could analyse in detail their connectivity patterns and their direct and indirect upstream and downstream partners (Figure 4A-O).

**Figure 4:**
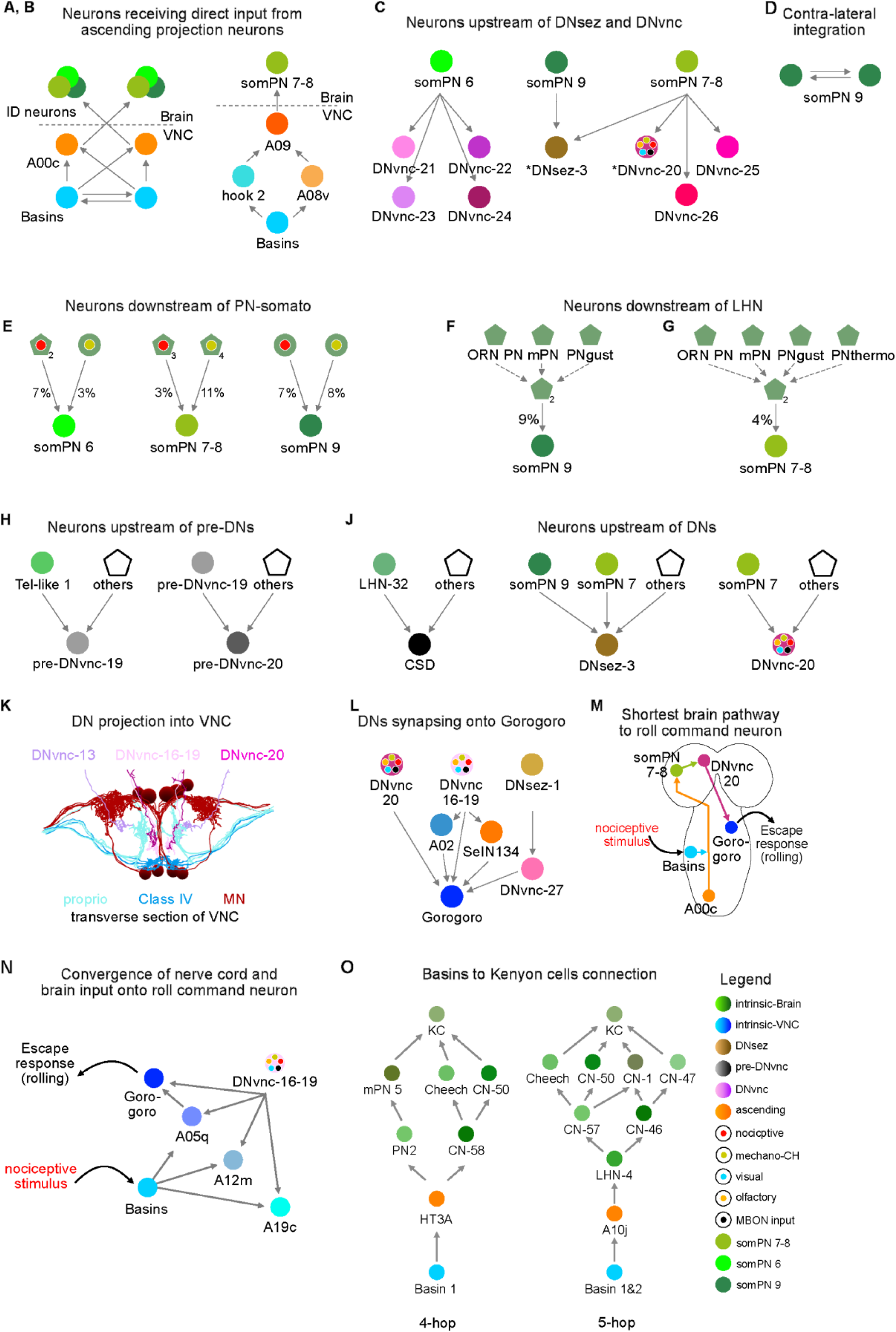
Circuit analysis. **A:** Shortest neuronal pathway from the VNC to the brain neurons somPN 6, somPN 7-8, and somPN 9 via the ascending neurons A00c. **B:** Path from the VNC to the brain via other ascending neurons. **C:** Brain neurons synapsing on DNs. *Identified neurons. **D:** Bilateral signal integration in the brain. **E:** 1-hop multimodal integration from PNsomato, and **F, G:** lateral horn neurons. Lateral horn neurons receive input from olfactory (ORN), gustatory (PNgust), and multiglomerular (mPN) projection neurons. The polygons indicate a group of neurons and the relative synaptic input. **H:** Brain neurons synapsing onto pre-DNvnc, and **J:** DNvnc neurons. **K:** DNvnc axon location in the VNC. **L:** DNvnc synapsing onto Gorogoro, the command neuron for rolling behaviour. **M:** Shortest brain pathway from Basins to Gorogoro via DNvnc-20 which also integrates olfactory, visual and MBON input **N:** DNvnc-16-19 integrate Basin, olfactory, visual and MBON input and synapse onto A05q and Gorogoro. **O:** Example 4-hop and 5-hop neuronal pathway from Basins to Kenyon cells. Synaptic threshold: ≥ 3 synapses.

Finally, to validate our methodology (Figure 5A-P), we searched for available GAL4 lines that selectively target gene expression to these neurons. We were able to identify lines for 3 out of the 26 neurons (Figure 5F, 5K and 5O). We used these GAL4 lines to selectively express GCaMP8s in each of these neurons, and imaged their responses to Basin activation using two-photon microscopy (Figure 1G-H and 5G, 5L and 5P). We confirmed that they all had similar responses when imaged individually, to the responses obtained using whole-brain imaging, thereby validating our ability to correctly identify neurons after whole-brain imaging of neural activity and confirming the reproducibility of their responses to noxious stimuli across individuals.

**Figure 5:**
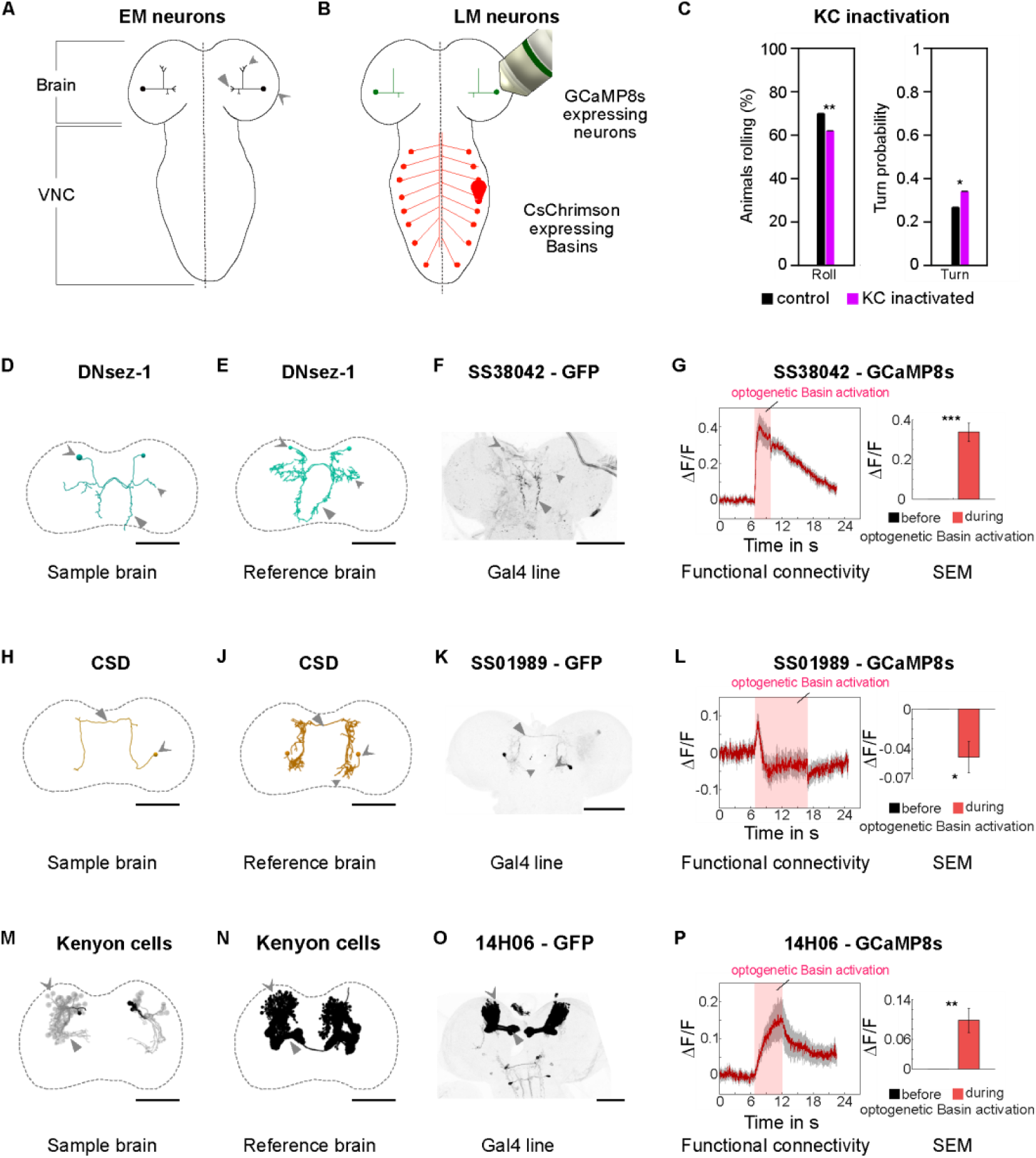
Neuron validation. **A:** Schematic of identified brain neurons. **B:** Schematic of the validation experiments in which GCAMP8s is selectively expressed in a specific neuron type (using a neuron-specific promoter) and its response to optogenetic activation (using 625 nm red LED illumination) of all Basins1-4 along the nerve cord (using 72F11 promoter to express Chrimson) is imaged using 2-photon microscopy. **C:** In animals with inactivated KCs [w;72F11-lexAp65; LexAop2-IVS-CsChrimson/UAS-A1ACR1-EYFP-p10/14H06-GAL4] (magenta), Basin activation (15 sec illumination with 625 nm red LED at 600 µW/cm²) evoked significantly less rolling (*P* = 0.0037), and more turning (*P* = 0.025) compared to controls [w;72F11-lexAp65; LexAop2-IVS-CsChrimson/ 14H06-GAL4] (black) within the Basin activation window. n = 366 larvae for the experimental group and n = 351 larvae for the control group; Chi-squared test. **D, H, M:** Reconstructed neurons in the sample brain. In **M**, the identified Kenyon cells are in black, and additional Kenyon cells are shown in grey. **E, J, N:** Corresponding neurons in the reference connectome. **F, K, O:** Matching driver lines expressing GFP. **G, L, P:** Neuronal responses to Basin activation recorded via 2-photon microscopy, and bar plot of mean and standard error of the mean (SEM) comparing neuronal activity before stimulus and during the complete stimulus window **(G, P)** or stimulus start +1s **(L)**. Paired-sample t-test. **G:** n = 8; *P* = 0.003. **L:** n = 8; *P* = 0.02. **P:** n = 11; *P* = 0.002. Scale bar: **D, E, H, J, M, N**: 25 µm; **F, K, O**: 100 µm. Dashed line indicates the midline, barbed arrowhead shows cell body, triangle indicates axon, and curved arrowhead shows dendrite. See also Figure S4.

### Identifying brain neuronal lineages activated by Basin activation

The *Drosophila* larval brain contains approximately 100 stereotyped neuronal lineages. Neurons from each lineage derive from a single neuroblast and follow a characteristic trajectory to the neuropil and enter the neuropil at a stereotypic location and project into similar neuropil areas ^46^. The lineage identity of each neuron can therefore be uniquely identified based on the trajectory its axon takes to the neuropil and the neuropil entry point.

To identify the lineage of each of the 119 neurons that significantly responded to Basin activation, we therefore manually traced their axons that lead from the cell body to the neuropil entry point and beyond (Figure 2D-Y). We were able to trace in this way 101/119 neurons (85%), and the remainder could not be traced due to image artifacts affecting some portions of the EM volume. Based on their neuropil entry point, we identified 25 neuronal lineages in which 1–30 neurons responded to Basin stimulation (Figure 2E-Y; Table S1).

We wondered whether neurons from lineages that received the most direct Basin input were more likely to respond to Basin activation, than others. Therefore, we analysed the published connectome of the *Drosophila* larval brain ^9^ and found that 82 lineages contain neurons that receive Basin input within ≤5 hops. 25 of these 82 lineages contained neurons that significantly responded to Basin activation in our whole-brain imaging experiment. Neurons within lineages connected to Basins by fewer than five hops responded to Basin stimulation (Table S2). Only 15 lineages contained neurons that received two-hop input from Basins (via direct input from ascending projection neurons from the nerve cord that in turn receive direct Basin input), but 10 out of these contained neurons that significantly responded to Basin activation (Figure 2E–Y; Table S1). Thus, 40% of the 25 lineages whose neurons responded to Basins received two-hop input from Basins. Overall, our data reveal that neurons from lineages that contain the most direct, two-hop, input from Basins are more likely to respond to Basin activation, than neurons from lineages that receive more indirect input via longer paths: 10/15 (67%) vs. 15/67 (22%).

Next we analysed the activity of the neurons within the same lineage. We discovered that all neurons within a single lineage responded similarly to the stimulus. The activity traces often overlapped, and the neurons within the same lineage were either all depolarised or all hyperpolarised upon Basin activation. The only exception was the cell lineage BLAv1/2, where 28/30 cells were depolarised and 2/30 were hyperpolarised (Figure 2G). The results indicate that the neurons within a lineage mainly respond in a similar fashion, either being depolarised, or hyperpolarised.

The mean and median number of neurons that responded to Basin activation per lineage, was small (5 and 2, respectively). Interestingly, a few lineages contained a large number of neurons that responded to Basins (Figure S1; Table S4): BLAv1/2 (30; 27% of all neurons in that lineage); BLVa3/4 (9 neurons; 9% of neurons in that lineage); BLVp1/p2 (12 neurons; 19% of neurons in that lineage) and BLD2-4 (17 neurons; 24% of all neurons in that lineage). These four lineages, therefore, contributed 57% of all Basin-responsive neurons in the brain (68/119, 57%). Significant fraction of neurons from these lineages also received two-hop input from Basins via direct input from ascending projection neurons from the nerve cord: 26%, 20%, 23% and 11% of neurons from BLAv1/2, BLVa3/4, BLVp1/p2 and BLD 2-4 (Table S5), respectively, and others receive three-hop input. Our findings therefore identify four lineages as major contributors of neurons that process escape-inducing somatosensory information in the brain.

### Identifying individual brain neurons responsive to Basin activation

Next, we randomly selected 50 neurons that significantly responded to Basin activation to determine what fraction of them we can identify as corresponding to one of the neurons from the reference brain. We traced their fine projections and compared their morphology to the reference neurons from our recently published reconstructions of all larval brain neurons ^9^. In our 12 x 12 x 12 nm/voxel resolution eFIB-SEM dataset, of the selected neurons, we were able to trace 18 cells sufficiently to identify them (DNsez-1, CSD, LHN-32, DNsez-3 left and right, somPN 9, MBON-q1, Noose 1, Tel-like 1, DNvnc-20 left and right, pre-DNvnc-20, DNvnc-13, and five Kenyon cells [KCs]); 8 neurons could be narrowed down to 2-4 candidates with very similar projections belonging to the same neuron types within the same lineage (DNvnc-14-15, somPN 7-8, DNvnc-16-19; two Lasso-top neurons that could each be one of Lasso-top 1-3, but since Lasso-top 1 only receives 7-hop input from Basins, whereas Lasso-top 2 and 3 both receive 5-hop input, it is likely that the identified Basin-responsive neurons are Lasso-top 2-3; and 3 neurons that each could be somPN 6, pre-DNvnc-18, or pre-DNvnc-19 [Figure 3E,F,H,L,N, and Table S2]). The remaining cells could not be traced sufficiently to allow identification. 20/26 identified neurons were depolarised and 6/26 hyperpolarised in response to Basin activation (Figure 3A,C-T, S2; Table S2).

All identified cells receive input from Basins via two to five hops (Table S2). Interestingly, we found that, as a population, the identified neurons that responded to Basin activation had significantly shorter paths in the connectome to Basins than all brain neurons (Figure S5).

Some of the identified neurons formed strong hubs (having ≥20 in- or out-degree) within the network, such as DNsez-3, DNsez-1, LHN-32, somPN 8, pre-DNvnc-20 and CSD (Table S2).

These neurons integrate information from multiple other neurons of the circuit. We also discovered recurrent feedback connections onto Basins from two of the hub neurons (LHN-32 and DNsez-1).

We analysed in more detail the 26 identified neurons (Table S2). Three identified brain neurons receive the shortest possible, two-hop, input from Basins, via direct synaptic input from ascending projection neurons (A00c) that receive direct Basin input: somPN 7-8, somPN 9, and somPN 6 (Figure 4A). SomPN 9 integrates information from the two sides of the body (Figure 4D). SomPN 7-8 also receives direct input from other somatosensory ascending projection neurons (Figure 4B and 4E). These three neurons also integrate Basin input with other sensory modalities (gustatory and olfactory via input from lateral horn neurons [LHNs], Figure 4F-G).

Furthermore, somPN 6, somPN 7-8 and somPN 9 directly innervate brain output neurons (DNs), thus participating in short nerve cord-brain loops (Figure 4C; Table S3). Thus, somPN 7-8 and somPN 9 directly innervate the DN-SEZ neuron DNsez-3. SomPN7-8 also directly innervates the identified DNvnc-20.

Overall, one third of the identified neurons that were responsive to Basins were DNs (19% DN-VNC and 12% DN-SEZ), transmitting the signal from the brain to the VNC, or to SEZ (Figure 4H, J, K). This includes five DN-VNC neurons (DNvnc-13, DNvnc-14 or DNvnc-15, DNvnc-16-19, DNvnc-20 left and right), and three DN-SEZ (DNsez-1, DNsez-3 left and right) (Table S2). Furthermore, four neurons were pre-descending neurons (pre-DN-VNCs), directly upstream of DNs (CSD, pre-DNvnc-18, pre-DNvnc-19, pre-DNvnc-20).

Two of the identified DNs that are activated by Basins synapse directly onto the nerve-cord command neuron for rolling, Gorogoro (DNvnc-16-19 and DNvnc-20, Figure 4L-M) ^13^ that also receives Basin input via shorter nerve cord pathways. DNvnc-16-19 also synapse onto A05q, a nerve cord interneuron presynaptic to Gorogoro that receives direct Basin input ^13^. Thus, even this shortest reflex pathway from Basins to Gorogoro is modulated by brain inputs. In addition to being activated by Basins, the brain DNs that synapse onto Gorogoro receive indirect input from other sensory modalities (olfactory and visual). This circuit architecture suggests that the strongest effect of other modalities on rolling could be in the context of Basin activation, when Basin input could facilitate the activation of DNs by other modalities.

We also found that some neurons previously implicated in processing olfactory stimuli are activated by Basins (a lateral horn neuron, LHN-32), suggesting these neurons are multisensory ^9^. Surprisingly, five of the intrinsic neurons (Kenyon Cells [KCs]) of the learning circuit (the mushroom body [MB]), also responded to Basin activation, as did an MB output neuron (MBON-q1). KCs are probably one of the most studied neurons in the *Drosophila* nervous system, but despite this, they were never previously reported to respond to noxious cues, as they are usually thought of as representing conditioned stimuli (CS), such as odours, temperature or light), but not unconditioned stimuli (US) that serve as punishments (such as noxious cues) or rewards during associative learning tasks. We identified a 5-hop synaptic pathway between Basins and KC (Figure 4O).

Finally, four neurons identified in this study as responsive to Basin activation, did not belong to any previously known neuron categories (Lasso-top 1-3 neurons, Tel-like 1, Noose 1).

Together, these findings expand our knowledge of nociceptive processing in the insect brain and identify numerous neurons for future follow-up studies.

### Validating identified brain neurons

Next, we wanted to validate our methodology and our findings by confirming that the specific neurons we identified as responding to Basin activation using whole-brain imaging of neural activity, are reproducibly activated by Basins. To do this, we wanted to selectively express a calcium indicator of neural activity in one identified neuron at a time and image their responses to Basin activation in multiple animals. We, therefore, looked for available GAL4 or LexA lines that drive expression, selectively, in the neurons that we identified as responding to Basin activation. We were able to find lines for 3/26 of the identified neurons: DNsez-1 and CSD, as well as a line that drives expression in all KCs ^47,48^.

We used these lines to express GCaMP8s, selectively in the neurons of interest, and the Basin-specific driver line to express the optogenetic activator of neural activity, Chrimson, selectively in Basin neurons ^13^. We optogenetically activated Basins and imaged the responses of KCs, DNsez-1, and CSD using two-photon microscopy. The results reproduced our findings from the whole brain imaging experiment, where Kenyon cells and DNsez-1 were depolarised in response to Basin activation, and CSD was briefly depolarised, and then strongly hyperpolarised throughout Basin activation (Figure 3C,D,O, 5A-B, D-P). We also checked whether there are any neurons in the neighbourhood of these neurons with similar responses and found that, while DN-SEZ-1 exhibits a robust, prolonged excitatory response to Basin activation, none of its 10 neighbours do (Figure S4). This further confirms that we identified the correct neuron and validated the only neuron that responded to Basins in that neighbourhood. For CSD, two of its neighbours were also inhibited but CSD has a very characteristic and distinct morphology from those neighbours (Figure S4). Thus, even if the two neighbouring neurons are also inhibited by CSD, our two-photon imaging experiments in the CSD neurons demonstrate that it is reliably, and reproducibly inhibited by Basins, in multiple individuals.

Overall, these findings validate our methodology and show we could reproduce findings from whole-brain, pan-neuronal imaging experiments by imaging activity of a single neuron type at a time, with cell-type-specific expression of GCaMP. Whole-brain imaging enabled us to identify many more neurons involved in processing escape-inducing somatosensory stimuli, for which selective driver lines currently do not exist. In the future, whole-brain imaging of neural activity will enable systematic characterisation of response properties of all neurons in the brain, to a range of stimuli and across a range of behavioural tasks.

### Kenyon cells contribute to innate behaviour

Perhaps one of the most unexpected findings in our study is that the intrinsic neurons of the MB, the KCs, known to be required for associative learning and memory and thought to represent CS, also respond to Basins whose stimulation serves as an unconditioned aversive reinforcement signal during associative learning tasks ^16,49^.

We, therefore, wondered whether KCs contribute to innate behaviour, in particular the innate escape response to noxious stimuli. To test this, we optogenetically silenced KCs [using the red-shifted inactivator of neural activity, Ruby ^50^], during optogenetic activation of Basin neurons [using the red-shifted activator of neural activity, Chrimson ^51^] and analyse larval rolling escape response, which is normally induced by Basin activation ^13^. We found that the probability of rolling was mildly, but significantly reduced, whereas the probability of turning was significantly increased, when KCs were silenced, compared to controls. Thus, silencing KCs causes inappropriate selection of turn, at the expense of the roll. This result supports a role for the MB in the appropriate selection of the innate escape response (Figure 5C).

This finding highlights further the value of an unbiased whole-brain imaging approach when identifying neuron types involved in specific functions.

### Comparing neural activations with connectome predictions

Having identified a brainwide network of neurons responsive to Basin activation *in vivo*, we next asked to what extent the known synaptic wiring of the brain predicts the observed calcium response properties (i.e. excitation vs. inhibition) of these identified neurons. To test this, we constructed a recurrent firing rate model of the larva brain with neuron connection strengths taken directly from the connectome by assuming strength is proportional to the number of synapses between two neurons. The signs of connections were fixed based on automatically predicted neurotransmitter profiles of neurons (predicted as in Eckstein et al. 2016 ^52^ based on Wang et al. 2024 ^53^. We then simulated the optogenetic activation of Basin neurons in silico and studied the effects of Basin activation on the simulated neuron firing rates of those neurons that we experimentally identified as responsive to Basin activation (Figure 6A). Specifically, we evaluated the model’s ability to correctly rank these neurons according to their experimentally observed response to Basins.

**Figure 6:**
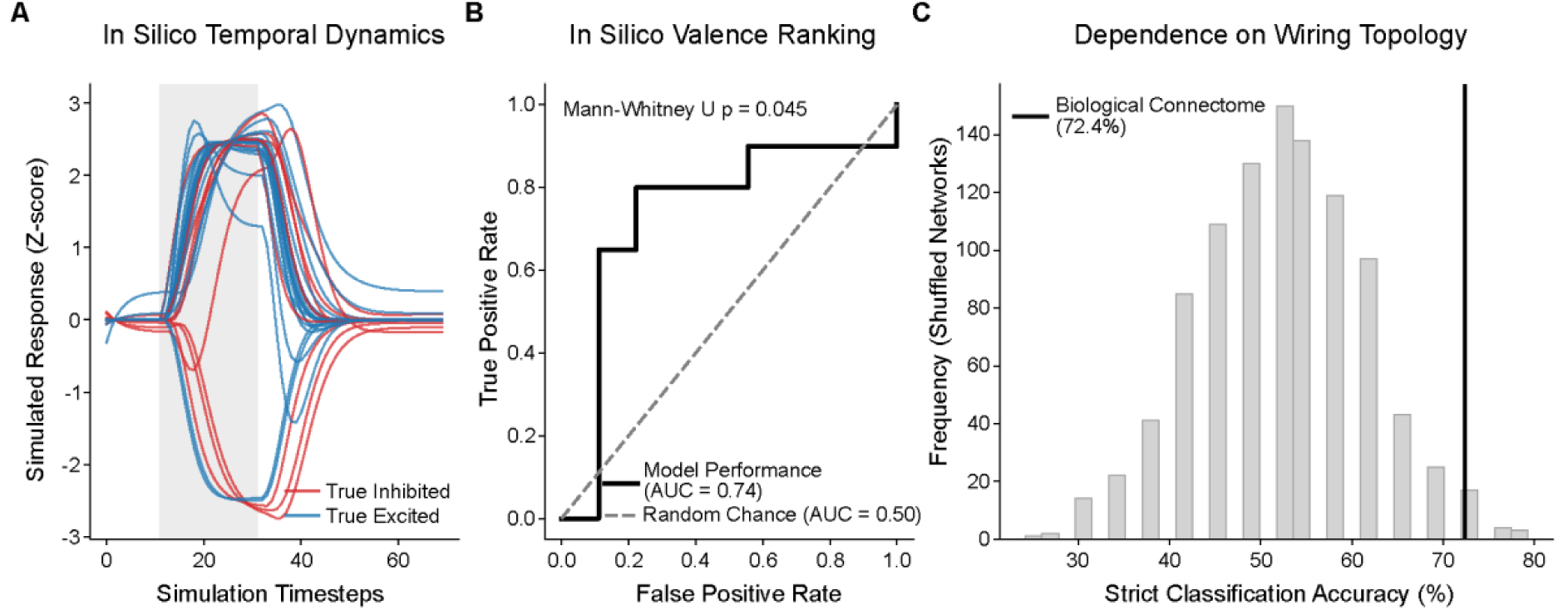
The static synaptic connectome predicts *in vivo* functional valence. (**A**) Z-scored temporal dynamics of individual *in silico* neurons during simulated Basin optogenetic stimulation (grey shaded region). Simulated firing rate traces are zero-centered to the pre-stimulation period and color-coded by their experimentally observed *in vivo* functional response (blue = true excited, red = true inhibited). (**B**) Receiver Operating Characteristic (ROC) curve evaluating the model’s ability to correctly rank these individual neurons by their observed *in vivo* valence. The connectome-constrained model (solid black line) successfully assigns higher simulated excitatory drive to experimentally excited neurons than to inhibited neurons, performing significantly better than random chance (dashed grey line; Area Under the Curve [AUC] = 0.74; Mann-Whitney U test, p = 0.045). (**C**) Permutation testing demonstrating the predictive dependency on the specific biological wiring topology. The strict classification accuracy of the biological connectome (72.4%, solid black vertical line) is significantly higher than a null distribution generated from 1,000 out-degree-preserving structurally shuffled networks (grey bars; Permutation test, p = 0.025). See also Figure S5, Table S5.

We found that the connectome-constrained model successfully segregated neurons based on their experimentally observed response sign and magnitude. *In silico* neurons corresponding to cells that were experimentally excited were driven to significantly higher simulated firing rates than those that were experimentally inhibited (Figure 6B). The model also accurately predicted the response sign (excitation > 0, inhibition < 0) for 72.4% of the evaluated neurons. To confirm that this predictive capability was driven by the specific biological architecture of the larval brain (i.e. rather than general network properties), we compared our model’s accuracy to 1,000 degree-preserving shuffled networks. The model with weights taken directly from the connectome yielded an accuracy significantly higher than the null distribution (Permutation test, *p = 0.025*) (Figure 6C). Overall, these findings suggest that the connectome predicts the responses of brain neurons to Basin activation, significantly better than would be expected by chance.

## DISCUSSION

In this study we developed a novel methodology for identifying neuron types and individual neurons following cellular-resolution, whole-brain imaging of neural activity with LSM. While cell body position alone is not sufficient to identify neurons, most neuron types (and even individual neurons) can be identified based on the locations and shapes of their projections. Our methodology, therefore, involves imaging the same brain with higher-resolution volume EM, to visualise neuronal projections, followed by reconstruction of the branches of those neurons that have interesting activity properties. This approach enables combining information about neuronal activity with previously available information about each neuron, such as their synaptic connectivity, molecular properties, or roles in behaviour.

We applied our approach to gaining a better understanding of the brain circuits downstream of somatosensory interneurons (Basins 1-4) that process the most threatening somatosensory (combination of noxious and mechanical) stimuli and that induce vigorous escape (rolling) and strong associative learning in *Drosophila* larvae.

After pan-neuronal calcium imaging in response to optogenetic activation of Basins 1-4 (in segment A4), we imaged the same CNS with eFIB-SEM at 12 x 12 x 12 nm/voxel resolution, sufficient to visualise medium-sized neuronal branches, and registered the two datasets of the same brain to each other, using landmark-based deformation method for registration (thin plate spline transformation). We quantified the registration error using leave-one-out cross-validation (LOOCV, see Methods for details) and found it is highly accurate (root mean square error across each dimension is X = 1.24 µm, Y = 1.86 µm, Z = 1.86 µm). Given that the width of a soma in the *Drosophila* larva is approximately 5 µm, these sub-cellular error margins confirm that the global transformation reliably and consistently maps EM coordinates to locations in light imaging space with cellular precision. One potential way of improving landmark-based registration, would be to use glial cells labelled with a fluorescent reporter of an additional colour (e.g. in glia in green if neurons are imaged with RGECO). Glial cells are visible in EM (without any label as they are darker), so they could be readily recognised in both imaging modalities and would add hundreds of landmarks.

From the LSM dataset we found that 119 neurons significantly responded to Basin activation. We randomly selected 50 of these neurons for follow up and manually traced their medium-sized branches in the eFIB-SEM volume. Because existing models for automated segmentation of neuronal arbours in *Drosophila* were trained on eFIB-SEM datasets that were imaged with 8 x 8 x 8 nm/voxel resolution ^45^, for this proof-of-principle study we manually traced neuron arbours and were, therefore, not able to follow-up all 119 neurons. 26/50 could be traced sufficiently to identify them, either uniquely, or to a neuron type with very similar projections within the same lineage. Finally, we validated our neuron identification using GAL4 lines to selectively drive GCaMP in some of the identified neurons and confirmed they reproducibly respond to optogenetic activation of Basins 1-4 in multiple individuals.

We also considered how resolution and imaging speed influence the applicability of this approach. Prior studies show that imaging *Drosophila* brains with eFIB-SEM using 8 x 8 x 8 nm/voxel resolution is sufficient to reconstruct neuronal fine branches and synapses ^8,45^. Here we chose a lower, 12 x 12 x 12 nm/voxel resolution, to assess the extent to which medium-sized branches will be traceable and the extent to which this would be sufficient for identifying neurons. While, for *Drosophila* larval brains, imaging with 8 x 8 x 8 nm/voxel resolution would be very fast (ten days per brain), imaging with 12 x 12 x 12 nm/voxel resolution is 30% faster, and would enable acquisition of large sample sizes, or faster imaging of larger brains. Overall, our study demonstrated that, after cellular-resolution imaging with LSM, imaging the same brain with 12 x 12 x 12 nm/pixel eFIB-SEM, enables identification of ca. 50% of neurons. In the future, for smaller brains, such as that of *Drosophila* larva, imaging with synaptic-resolution (8 x 8 x 8 nm/voxel) would be fast enough and would provide significant advantages: first, it would enable tracing of fine branches and synapses; second, it would enable automated segmentation of neuronal branches using existing models and therefore identification of most neurons in the brain ^45^. As EM imaging and reconstruction gets faster and faster, it may be possible to apply this approach to larger and larger brains ^30–32,54^. Furthermore, expansion microscopy may also be used instead of EM to visualise neuronal projections following whole-brain imaging of neural activity ^55–58^.

Somatosensory circuits in the *Drosophila* larval nerve cord have been extensively studied, but much less was known about somatosensory signalling in the brain. Here we discovered multiple novel neuron types responsive to Basin activation. 20/26 (77%) of identified neurons were depolarized and 6/26 (23%) hyperpolarized in response to Basin activation. We discovered hub-neurons for signal processing, as well as feedback neurons that signal back to Basins, likely modulating their response to sensory stimuli. Identified neurons also included direct targets of ascending projection neurons in the brain, some of which integrate information from two sides of the body, while others integrate somatosensory information with other sensory modalities, as well as brain descending neurons that project to the premotor regions of the nerve cord and could play a role in facilitating escape.

Surprisingly, we also discovered Basin responses in neuron types and brain areas previously associated with olfactory processing and learnt responses to conditioned stimuli. For example, a subset of KC, the intrinsic neurons of the learning circuit (the MB), responded to Basin activation. While Basin 1-4 activation was previously shown to serve as an unconditioned stimulus and drive strong associative learning when paired with odour by activating dopamine neurons^15^, it was not known to activate KCs. KCs have been demonstrated to encode conditioned stimuli, such as odours, light, or temperature that can be associated with unconditioned stimuli, such as noxious stimuli ^59–63^. Our study shows that a subset of KCs responds to Basin stimulation. Indeed, analysing the connectome reveals axo-dendritic paths from Basin neurons to a subset of KCs. Furthermore, we found that optogenetic silencing of KCs reduces the likelihood of innate rolling escape response to nociceptive interneuron activation. This suggests that KCs mediate not only learnt behaviour but also facilitate innate escape response. In the future, it will be interesting to test whether the KCs activated by noxious stimuli also play a role in learning to respond more or less strongly to noxious stimuli depending on experience.

Overall, these findings demonstrate how unbiased, whole-brain imaging approaches can reveal unexpected response properties, even for some of the most studied cell types in the brain. Furthermore, for many of the identified neurons, GAL4 lines are not available, underscoring the importance of whole-brain imaging to study the response properties of a large fraction of neurons for which genetic driver lines are not available.

In summary, our study identified multiple neuron types in the insect brain involved in processing threatening, escape-inducing somatosensory stimuli. The methodology we have developed will enable overlaying brainwide activity maps during various behavioural tasks onto connectivity maps, an essential step towards understanding neural circuit function.

## RESOURCE AVAILABILITY

### Lead contact

Requests for further information and resources should be directed to and will be fulfilled by the lead contact, Marta Zlatic.

### Materials availability

This study did not generate new unique reagents, and plasmids. Generated transgenic flylines are available upon request from lead contact.

### Data and code availability

#### Data

● Electron microscopy and light-sheet microscopy calcium traces dataset has been deposited at VirtualFlyBrain: https://virtualflybrain.org/about/hosted/
● Behaviour data and 2-photon imaging data reported in the paper will be shared by the lead contact upon request.

#### Code

All original code has been deposited at [Zenodo/ Github] and is publicly available at: https://github.com/ZlaticLab/Randel-et-al.

#### Additional information

Any additional information required to reanalyze the data reported in this paper is available from the lead contact upon request.

## ACKNOWLEDGEMENTS

The authors thank MRC Laboratory of Molecular Biology, HHMI Janelia Research Campus and Department of Zoology, University of Cambridge for funding and support. We thank Bill Lemon (HHMI Janelia Research Campus), Fly core (especially Monti Mercer) at JRC, and Nan Hu and Oxana Elliott at the Department of Zoology, University of Cambridge for fly crosses. We thank the Janelia Visitor Project for outstanding support over the years. We thank MRC Laboratory of Molecular Biology Core funding (MZ, AC, MJW and KW), HHMI Janelia Research Campus (MZ, AC, PJK, NR, CW, HFH, SCX, SP), European Union Horizon 2020, Marie Skłodowska-Curie Grant No 838225 (NR), Wellcome Trust grant 205038/Z/16/Z (AC), Wellcome Trust grant 205050/Z/16/Z (MZ), ERC grant ERC-2018-COG: 819650 (MZ, NR, ASC) for funding.

## AUTHOR CONTRIBUTIONS

NR: Conceptualisation, Formal analysis, Investigation, Methodology, Validation, Project administration, Visualization, Writing – original draft, Writing – review and editing, Funding acquisition. CW: Methodology. MSC: Data curation. Software. KW: Validation. Methodology. Investigation. SP: Resources. CSX: Resources. AC: Data curation. HH: Conceptualisation. Funding acquisition. Resources. AC: Data curation. Funding acquisition. PJK: Conceptualisation, Software, Project administration, Data curation. Resources, Funding acquisition. MZ: Conceptualisation. Project administration, Resources, Writing – original draft, Writing – review and editing, Supervision, Funding acquisition.

## SUPPLEMENTAL INFORMATION

Document S1. Figure S1-S5, Table S1-S5, Standalone files, and supplementary reference.

## STAR★METHODS

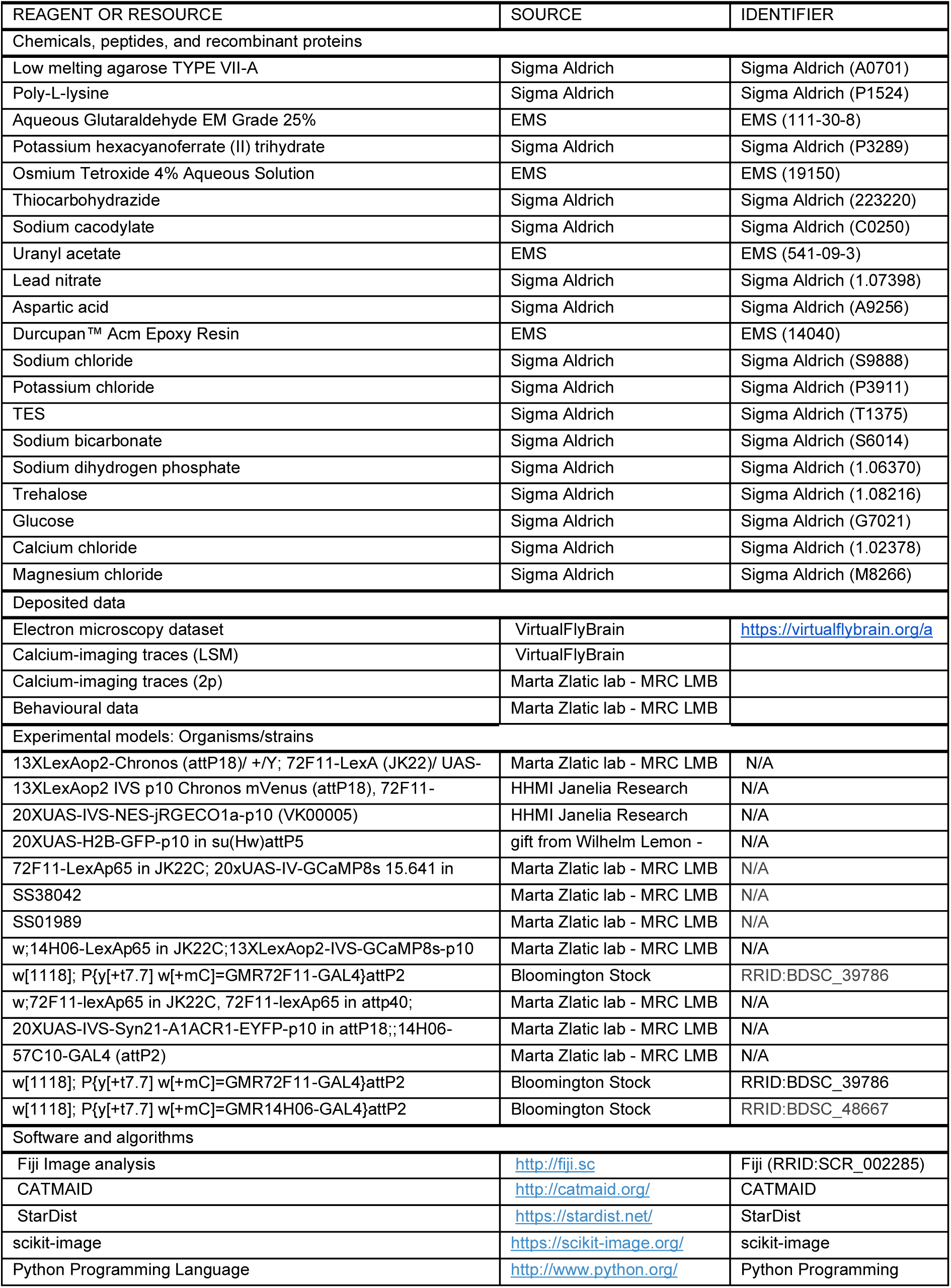
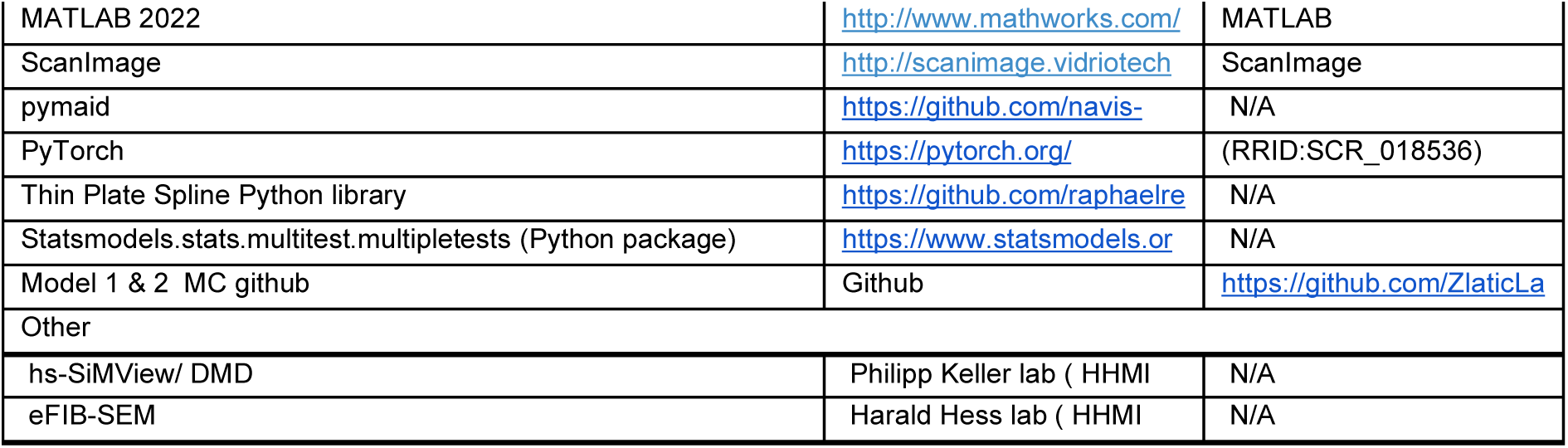
KEY RESOURCE TABLE.

## EXPERIMENTAL MODEL AND STUDY PARTICIPANT DETAILS

### Organisms/fly strains

#### Whole-brain calcium imaging

The experiment was conducted using late first-instar *Drosophila melanogaster* larvae. Flies expressed UAS-RGECO1a pan-neuronally for monitoring neuronal activity, and LexAop-Chronos in all Basins (72F11-LexAp65), for optogenetic activation ^13,51,64,65^.

The final cross 13XLexAop2-Chronos (attP18)/ +/Y; 72F11-LexA (JK22)/ UAS-H2B-GFP (attP5); UAS-RGECO1a (VK0005)/ 57C10-GAL4 (attP2) was made by using the following stocks: 13XLexAop2 IVS p10 Chronos mVenus (attP18), 72F11-LexAp65 (JK22C), and 20XUAS-IVS-NES-jRGECO1a-p10 (VK00005) from public collection at HHMI Janelia Research Campus (Ashburn, VA, US); 20XUAS-H2B-GFP-p10 in su(Hw)attP5 gift from Wilhelm Lemon - Philipp Keller lab, HHMI Janelia Research Campus (Ashburn, VA, US); and 57C10-GAL4 (attP2) from Marta Zlatic lab - MRC LMB (Cambridge, UK) ^13,37,65^.

#### Two-photon imaging

The experiment was conducted using third-instar *Drosophila melanogaster* larvae. Neuronal activity was recorded from specific cells expressing GCaMP8s during optogenetic activation of Basins that express CsChrimson ^51,66,67^.

Crosses were made using the following stocks: 72F11-LexAp65 in JK22C; 20xUAS-IV-GCaMP8s 15.641 in attp2, 13XLexop2-CsChrimson-tdTomato in VK00005 crossed with the GAL4 for DNsez-1 (SS38042) and CSD (SS01989), and Kenyon cells ((w;14H06-LexAp65 in JK22C;13XLexAop2-IVS-GCaMP8s-p10 50.641 in VK00005, 20XUAS-CsChrimson-mCherry-trafficked in su(Hw)attP1 with w[1118]; P{y[+t7.7] w[+mC]=GMR72F11-GAL4}attP2 ^37,65^).

#### Behavioural experiments

The experiment was conducted using third-instar *Drosophila melanogaster* larvae. Larval behaviour was recorded with a custom-made rig before and during optogenetic activation of Basins and/or Kenyon cells ^33^. The cross of w;72F11-lexAp65 in JK22C, 72F11-lexAp65 in attp40; 13XLexAop2-IVS-CsChrimson.tdTomato in VK00005/TM6B to 20XUAS-IVS-Syn21-A1ACR1-EYFP-p10 in attP18;;14H06-GAL4 in attP2/TM6B was used as the experimental group while the control group to w[1118]; P{y[+t7.7] w[+mC]=GMR14H06-GAL4}attP2 ^50^.

## METHOD DETAILS

### Light-sheet microscopy

#### Sample preparation

Flies were raised on standard fly food. For the experiments, eggs were incubated at 25°C on standard fly food with 0.5 mmol/l retinal. First instar larvae were dissected in physiological saline solution ^68^, and embedded in 1% low-melting temperature agarose (Sigma Aldrich Low melting agarose TYPE VII-A, Sigma (A0701)) in the physiological saline solution, as described in ^26^.

#### Data acquisition and processing

The sample was imaged using a high-speed multi–view (hs-SiMView) light–sheet microscope at HHMI Janelia Research Campus ^26,69^. Volumetric imaging was performed at volume rate of 2.87 Hz and with a voxel size of 0.406 × 0.406 × 1.7 µm. To reduce motion and deformation artefacts, we imaged activity in the extracted nervous system. To account for any sample drift during activity imaging, we took the first volume in the time series as our static reference and used phase cross-correlation to calculate the rigid shift for every subsequent volume and then estimate the shift across the entire time series.

A 561 nm laser was used for fluorescence excitation for the purpose of whole-brain imaging, while a 488 nm laser coupled into one of the microscope’s detection arms using a dichroic mirror was concurrently used for optogenetic activation of all Basin neurons (Basin-1, -2, -3 and -4, i.e. four pairs) in abdominal segment four.

The patterned manipulation arm was implemented in a high-speed simultaneous multi-view light sheet microscope by projecting a laser beam onto a digital micromirror device (DMD) and relaying the DMD pattern into the microscope through a 1:1 relay-lens pair, dichroic mirror, 150 mm tube lens, and a Nikon 16× water-dipping objective (focal length 12.5mm). The DMD pattern was demagnified by approximately 12× at the sample plane, corresponding to an effective projected DMD pixel size of approximately 0.9 µm. The projected DMD field covered approximately 830 µm at the sample plane, enabling wide-field spatial patterning of the manipulation light. Before each experiment, the illuminated DMD region was projected onto the objective/image plane and spatially registered to ensure accurate mapping between the DMD pattern and the sample field of view. The integrated optical and control setup enabled flexible spatiotemporal optogenetic stimulation by allowing user-defined patterns to be projected, switched on and off, flashed, and repeated with programmable pulse duration, stimulation frequency, number of repetitions, and total stimulation duration. This provided a comprehensive manipulation platform in which the spatial distribution and temporal sequence of stimulation light could be precisely coordinated with live imaging.

The optogenetic activation protocol alternated between 2 s and 5 s periods of optogenetic stimulation, with a 200 s pause between successive stimulation periods (n = 18). The resulting datasets were processed using a custom analysis pipeline as previously described ^26,69^.

#### Electron microscopy sample preparation and imaging

After functional imaging, the sample was immediately prepared for EM volume acquisition (sample ID: 1099). It was fixed in 4% glutaraldehyde in 0.1 M sodium cacodylate buffer (pH 7.4) for 1h at 4°C, and left in the agarose cylinder during the whole preparation. All washing steps were carried out with 0.1 M sodium cacodylate buffer, until the thiocarbohydrazide step, when water was used.

Post-fixation was performed with 1.25% potassium hexacyanoferrate (II) trihydrate, 1% OsO_4_ in 0.1 M sodium cacodylate buffer for 30 min at 4°C. Next, the sample was incubated for 10 minutes in 1% filtered thiocarbohydrazide in water at room temperature, followed by 1% uranyl acetate in water overnight at 4°C. Afterwards, the sample was incubated in pre-warmed Walton’s lead aspartate for 1 h at 60°C ^70^. Finally, the sample was incubated in 0.8% OsO_4_ for 30 min on ice. Following that step, the sample was dehydrated in acetone and embedded in Durcupan resin (EMS14040). The Durcupan block was finally mounted on customised soft-copper studs, which are required for the modified eFIB-SEM pipeline ^32^. After trimming and coating with 10 nm gold and 100 nm carbon, the whole sample was imaged at 12 x 12 x 12 nm/voxel.

#### Landmark-based registration of EM and LSM volume

After light-sheet imaging of neural activity, fixing and preparing the sample for EM imaging introduces a shrinking in size and non-linear deformations. We therefore use a non-rigid, landmark-based deformation method for registration, the thin plate spline transformation (TPS). Using 51 landmarks that are evenly spaced throughout the brain, ensures that non-linear deformations are taken into account and the method is robust.

To translate points between EM and LSM coordinate spaces, we identified 51 shared landmarks in the brain lobes using Fiji/BigWarp (RRID:SCR_002285) ^71^. The landmarks were selected, according to cell morphology and arrangement based on the moving and fixed points from the EM and LSM spaces, respectively. We calculated a TPS using the Thin Plate Spline Python library (https://github.com/raphaelreme/tps) with zero regularization (α = 0.0) with Python Programming Language (RRID:SCR_008394).

Our quantification of registration error using leave-one-out cross-validation (LOOCV) demonstrates that this registration is highly accurate (*root mean square error across each dimension is X = 1.24 µm, Y = 1.86 µm, Z = 1.86 µm)*.

#### Validation of EM-LSM registration

To evaluate the accuracy of the transformation between the LSM and EM volumes, we quantified the registration error utilizing leave-one-out cross-validation (LOOCV) ^72^. To ensure geometrically accurate deformation, all landmark coordinates were first mapped into an isotropic physical space (µm). During each iteration of the LOOCV, a single landmark pair was withheld to act as unseen test data, and the thin-plate-spline (TPS) model was fitted to the remaining landmarks. This model was then used to predict the LSM location of the withheld EM coordinate. Registration accuracy was quantified by calculating the 3D Euclidean distance (offset vector) between the transformation-predicted location and the true manually-defined location of the withheld landmark.

Evaluation of the transformation via LOOCV demonstrated a reliable global registration. The median target registration error across the volume was 1.54 µm, with an overall mean absolute error (MAE) of 2.29 µm. The overall root mean square error (RMSE) for the volume was 2.91 µm. Breaking down the error by spatial axis revealed tightly constrained variance in RMSE results across each dimension of space (X = 1.24 µm, Y = 1.86 µm, Z = 1.86 µm). Given that the width of a soma in the *Drosophila* larva is approximately 5 µm, these sub-cellular error margins confirm that the global transformation reliably and consistently maps EM coordinates to locations in light imaging space with cellular precision.

#### Soma segmentation

Somas in the EM volume were segmented using a convolutional neural network [StarDist (RRID:SCR_027715)] ^73^ and trained using a manually annotated dataset of raw images and soma segmentation masks. This process identified 11,596 putative cells across the imaged sample. The centroid of each EM soma was then determined using scikit-image (RRID:SCR_021142) functions ^74^, and projected into the light-sheet volume using the previously described TPS transformation. The LSM-projected centroids were then used to generate soma segmentations in the light-sheet volume. To do this, a 3μm^3^ sphere mask (corresponding to the average soma size in larval *Drosophila*) was defined and positioned at the central location of every soma. Once applied to every cell, this process generated an instance segmentation of every soma in the LSM volume, based on the initial EM segmentation.

#### Extraction of neuronal activity

To extract the calcium fluorescence for each soma, the LSM data was first spatially filtered to reduce low-spatial frequencies in each volume, and therefore reduce signal blur between neighbouring cells. To do this, unsharp masking was applied to each LSM volume in time using the unsharp_mask function in scikit-image (radius=5.0, amount=6.0). For each cell, the mean calcium fluorescence was extracted by calculating the mean intensity across all voxels belonging to the cell for each point in time. This generated a time-series of spatially-filtered fluorescence values for each cell. Further analyses focused only on somas in the brain and supraesophageal zone (SEZ). To do this, a bounding box was drawn around these regions and all somas positioned outside of this box were excluded (i.e. those belonging to cells in the nerve cord). This process resulted in a total of 3,480 remaining neurons.

To study soma calcium responses to Basin stimulation, we calculated stimulus-evoked calcium traces for every cell. For each stimulus delivery (n=18), fluorescence values were taken from ∼5 s before and ∼5 s after stimulus onset (pre-window=15 frames; post-window=15 frames; imaging frequency=2.87 Hz). For every cell, stimulus-evoked fluorescence values were normalised to the pre-stimulus baseline using the following equation:

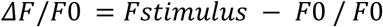

F_stimulus_ is the mean fluorescence value during stimulus delivery, while F_0_ is the mean calcium signal during the ∼5 s before stimulus delivery. For every cell and stimulus delivery, the average fluorescence throughout the stimulus-delivery period was calculated (called ‘stimulus responses’ from here on).

#### Neuron identification

Somas showing statistically significant responses to stimulation were located in the EM volume of the sample brain. The axons of selected somas were manually reconstructed using CATMAID (RRID:SCR_00627) ^75^. To identify the cells, neurons of the same lineage were reconstructed until lineage identity could be established. By comparing lineages and neurons across the sample and reference brain, cell lineage and single-cell identity was determined.

#### Reconstruction of neurons in eFIB-SEM volume

Neurons in the registered 12 x 12 x 12 nm/pixel resolution eFIB-SEM volume were manually reconstructed using CATMAID, as previously described ^9,75^.

#### Synaptic connectivity and circuit analysis of identified neurons

Once neurons were reconstructed in the eFIB-SEM volume and identified based on their morphological similarity to neurons in the previously published reference volume ^9^. The synaptic connectivity and circuitry of identified neurons was analysed in the reference volume, using pymaid (https://github.com/navis-org/pymaid) with axon-dendritic connections (threshold: 0.01), as previously described ^9^.

#### Analysing paths from Basins to downstream neurons

To identify the fraction of neurons that are a minimum of 2-, 3-, 4-, or 5-hops downstream of Basins, we performed network analysis using the published brain ^9^ and nerve-cord connectivity matrices ^13^. Only axo-dendritic connections were used and the connectivity matrices were thresholded to retain only connections comprising at least three synapses (since numerically weaker connections are not reproducible and could be due to developmental noise ^76^. To measure the propagation of signals from Basin neurons to brain neurons, we simulated stochastic network cascades as previously described ^9^. In short, synaptic transmission through the network was simulated across 100 independent runs, with the probability of signal propagation between two neurons being drawn directly from the synaptic strength between those neurons. For each brain neuron, we counted the number of times it was visited after 1-, 2-, 3-, 4- or 5-hops. To eliminate low-probability stochastic noise, neurons had to be visited more than once during these simulations to be considered to be connected via that number of hops. Neurons that failed to receive a signal within the maximum 5-hop threshold were excluded from the distance analysis.

### Neuron validation

#### Larval Dissection and Mounting

Central nervous systems (CNSs) of third instar *Drosophila* larvae were dissected in cold saline composed of 103 mM NaCl, 3 mM KCl, 5 mM TES, 26 mM NaHCO₃, 1 mM NaH₂PO₄, 8 mM trehalose, 10 mM glucose, 2 mM CaCl₂, and 4 mM MgCl₂, with an osmolarity of 285 mOsm/L and pH ∼7.2. Dissected brains were mounted onto a small piece of cover glass coated with poly-L-lysine (P1524; Sigma) and placed in a 35 mm petri dish lid (Falcon Easy Grip Style; Code: 351008; Scientific Laboratory Supplies), with the brain hemispheres facing upwards.

#### Two-Photon Imaging and Optogenetic Stimulation

A mode-locked Ti:Sapphire femtosecond pulsed laser (Spectra-Physics), tuned to 920 nm, was used for GCaMP8s imaging. Fluorescence was detected with two photomultiplier tubes (Hamamatsu) following band-pass filtering. Larval brains were imaged at 44 frames per second (fps) in a single focal plane using a two-photon microscope equipped with a fast resonant/galvo scan module (Bergamo, Thorlabs), controlled via ScanImage (RRID:SCR_014307) 2022.

Optogenetic stimulation was delivered via a high-power red LED (4-Wavelength High-Power LED Source; Thorlabs), illuminating neurons through the main optical path of the Bergamo microscope and a 25× water immersion objective (NA 1.1, WD 2 mm; Nikon). Light stimuli were delivered for 10 s (CSD), 3 s (DNsez-1), and 5 s (Kenyon cells) with an interstimulus interval of 20 s (CSD), 27 s (DNsez-1), and 15 s (Kenyon cells). Each trial consisted of four stimulations, automatically triggered by ScanImage 2022.

#### Image Analysis

Image time series were processed in Fiji and further analysed using custom scripts in MATLAB R2022a (RRID:SCR_001622). Regions of interest (ROIs) were manually defined based on the standard deviation projection of the entire time series. A background ROI was also selected outside the target cell region in each dataset to control for LED light bleed-through into the PMT for the green channel. Fluorescence signals in target ROIs were corrected by subtracting this background signal.

Fluorescence changes (ΔF/F₀) were calculated as ΔF/F₀ = (Fₜ − F₀)/F₀, where F₀ is the mean fluorescence over a 2-second period preceding the onset of stimulation, and Fₜ is the mean fluorescence of the ROI at time t. For each brain, mean ΔF/F₀ was computed across the four consecutive stimulations and averaged over three trials.

#### Behavioural experiments

*Drosophila* larvae were tested using the laboratory’s custom-designed large-scale behavioral rig for optogenetic activation, as previously described ^33^. For each recording, approximately 40 early third instar larvae were transferred onto a 3% agar plate using a fine paintbrush. Larvae were allowed to crawl freely for ∼15 s across the plate prior to recording.

Behavioral responses were assessed during simultaneous optogenetic activation of Basin neurons and optogenetic inactivation of Kenyon cells. In the optogenetic protocol, 625 nm red LED illumination (600 µW/cm²) was initiated 30 s after the start of the recording and maintained for 15 s, followed by a 30 s light-off period.

Rolling behavior was detected as previously described and the percentage of animals rolling within the Basin activation window was quantified ^14,77,78^.

#### Connectome-constrained firing rate model

To investigate the mechanistic relationship between the static structural connectome and the functional responses observed during whole-brain imaging, we constructed a recurrent firing rate model constrained by the synaptic wiring of the *Drosophila* larval brain.

#### Connectome-contrained network generation

The network architecture was defined by the adjacency matrix extracted from the EM connectome. To account for variations in total synaptic output, the synaptic weights were normalized per presynaptic neuron (such that the sum of squared weights for each outgoing row equaled 1) and globally scaled by a factor of 0.5. This normalization was applied to ensure that firing rates did not explode to unphysiological levels during Basin stimulation. Neurotransmitter identities were incorporated by assigning negative synaptic signs (−1) to neurons predicted to be GABAergic or glutamatergic from EM imagery. All other neurons were modeled as excitatory (+1). This resulted in an effective weight matrix W_eff_, containing 3,460 neurons.

#### Neuron dynamics and Basin activation

The firing rate vector v(t) for every neuron in the network was simulated using a discrete-time leaky integrator model:

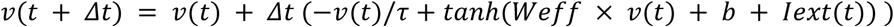

Where *Δ* is the simulation time step (0.5), *τ* is the timescale constant (1.0), b is a baseline activity vector (0.1), and I_ext_(t) represents external stimulus currents. The hyperbolic tangent function (tanh) was used to introduce non-linear saturation to the firing rates. To simulate the optogenetic activation of the Basin circuit, an external step current (I_ext_ = 10) was injected into the *in silico* Basin neurons (types 1-4, bilateral) for 20 timesteps. The model was implemented and simulated using PyTorch (RRID:SCR_018536).

#### Comparison of observed and modelled responses

Because basic linear firing rate models lack the biological complexities (e.g. gap junctions, precisely mapped neuromodulator tones, and non-linear dendritic integration) that needed to reliably simulate absolute membrane hyperpolarization, we evaluated the model based on its ability to accurately rank the relative valence of the evoked responses. The simulated evoked response for each neuron of interest was quantified by averaging its firing rate during the stimulation window and subtracting its pre-stimulation baseline firing rate. We then performed a Receiver Operating Characteristic (ROC) analysis to quantify the model’s ability to assign higher excitatory drive to *in vivo* excited neurons compared to *in vivo* inhibited neurons, calculating the Area Under the Curve (AUC) and using a Mann-Whitney U test.

To rigorously assess whether the connectome’s predictive power was significantly dependent on its specific biological topology, we generated 1,000 structurally shuffled network models using an out-degree-preserving shuffle. Specifically, the post-synaptic targets within each presynaptic neuron’s row in the weight matrix were randomised, destroying the specific biological topology while preserving the exact out-degree distribution and weight magnitudes of every neuron. The classification accuracy of the biological model (defined as the percentage of correctly predicted response signs) was compared against the null distribution of accuracies from all 1,000 shuffled models in order to calculate a permutation p-value.

## QUANTIFICATION AND STATISTICAL ANALYSIS

### Statistical analysis for whole-brain functional imaging

The reliability of cell responses to Basin stimulations were assessed by performing a one-samples t-test on the distribution of stimulus responses for each cell, assessing whether the sample mean differed significantly from 0. Given the large number of neurons, and therefore statistical tests, p-values were corrected for false discovery rate using the Benjamini–Hochberg (negative) procedure ^79^ (alpha = 0.1; implemented in the Python package, statsmodels.stats.multitest.multipletests (https://www.statsmodels.org/dev/generated/statsmodels.stats.multitest.multipletests.html)). A total of 119 cells had p-values lower than the adjusted significance threshold, and were therefore identified as responding neurons to be brought forward for further analyses.

### Statistical analysis for neuron validation

Differences in mean fluorescent signals (ΔF/F) between 7 s before and during the optogenetic stimulation were analyzed using a paired-sample t-test. A two-tailed α level of 0.05 was applied. All analyses were conducted in MATLAB R2022a. The N and actual P - values are reported in the Figure legends.

### Behavioural classification and statistical analysis

Larval behaviours were classified into six distinct categories based on standard locomotor phenotypes: crawl, turn, hunch, back-up, roll, and stop, using our previously published automated behaviour detection pipeline^14^.

For highly frequent behaviours (crawl and turn), statistical analysis was performed on a per-timestamp basis to capture temporal probability. For each video frame/timestamp within the window, the presence of the behaviour was scored as a positive event. The behavioural probability was defined as the proportion of positive timestamps relative to the total number of timestamps within the window for all the animals. The experimental and control groups comprised n = 366 and n = 351 larvae, respectively and were compared using a chi-squared test. To perform a Chi-square test of independence between the experimental and control groups, these probabilities for each group were converted to categorical counts by multiplying the calculated probability by the total number of larvae in each respective group.

For the remaining rare behavioural categories (e.g. roll), analysis was based on the overall fraction of responding individuals within a Basin activation window. A larva was scored as positive if it exhibited the target behavior at least once during the activation window. The response fraction was calculated as the number of positive larvae divided by the total number of larvae in the group. These categorical counts were then used to construct a contingency table for Chi-square analysis comparing the experimental and control groups.

For all analyses, p-values were adjusted for multiple comparisons across behavioural categories using the Benjamini–Hochberg false discovery rate (FDR) correction (α=0.1) for both the High-Frequency and High-Frequency behaviours, respectively ^79^.

