## Supplemental Information for "Combining brain-wide activity imaging with electron microscopy reveals a distributed brain network for processing threatening somatosensory stimuli"

### Supplementary Information Legends

#### Figures

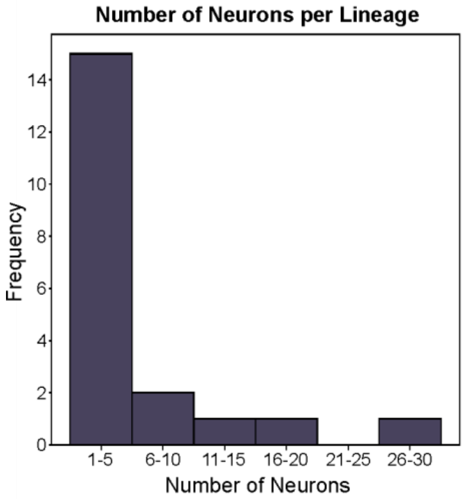

**Figure S1:** Histogram of the number of neurons per lineage that respond to Basin activation, related to Figure 2.

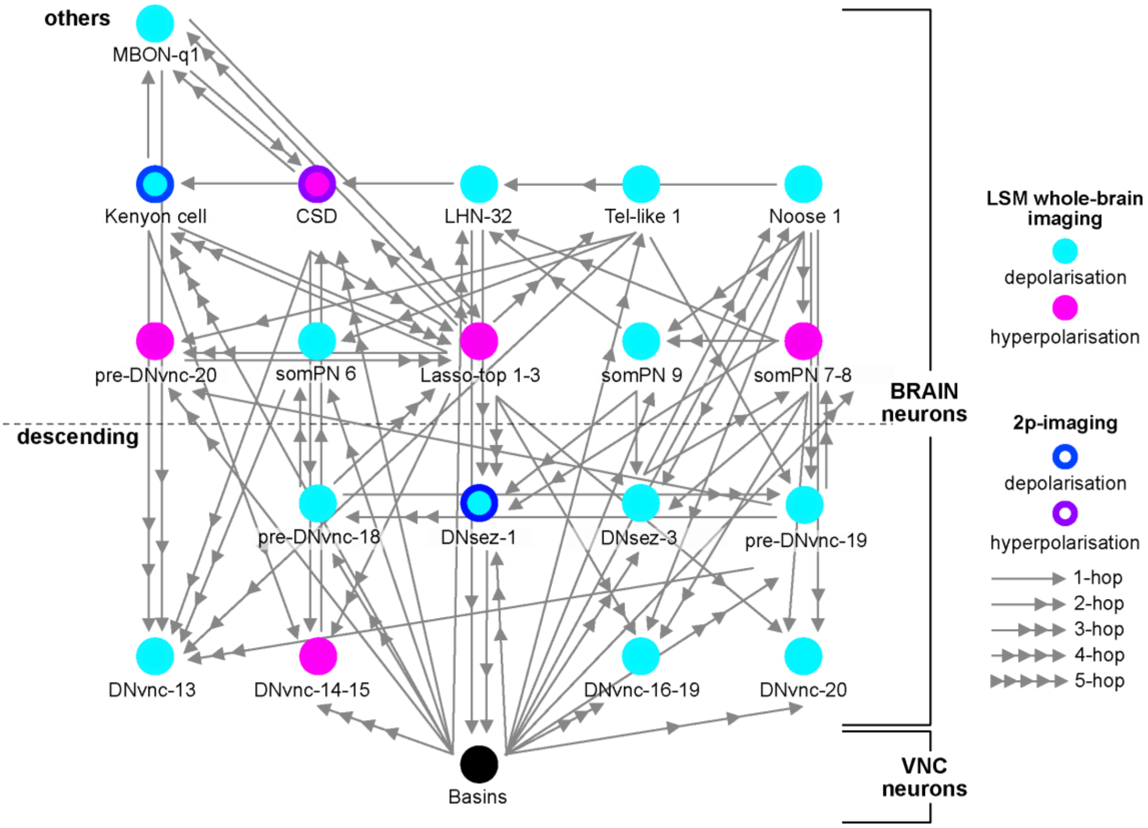

**Figure S2:** Synaptic connectivity of identified neurons that responded to Basin activation, related to Figure 3. Shortest axon-dendritic connections for all identified neurons within ≤4hops downstream of Basins (synaptic threshold: 0.01), and 5-hops connection from Basins to KC (synaptic threshold: ≥3 synapses). The number of arrowheads indicates the number of hops. Depolarised neurons are shown in cyan/blue, and hyperpolarising neurons are shown in magenta/violet.

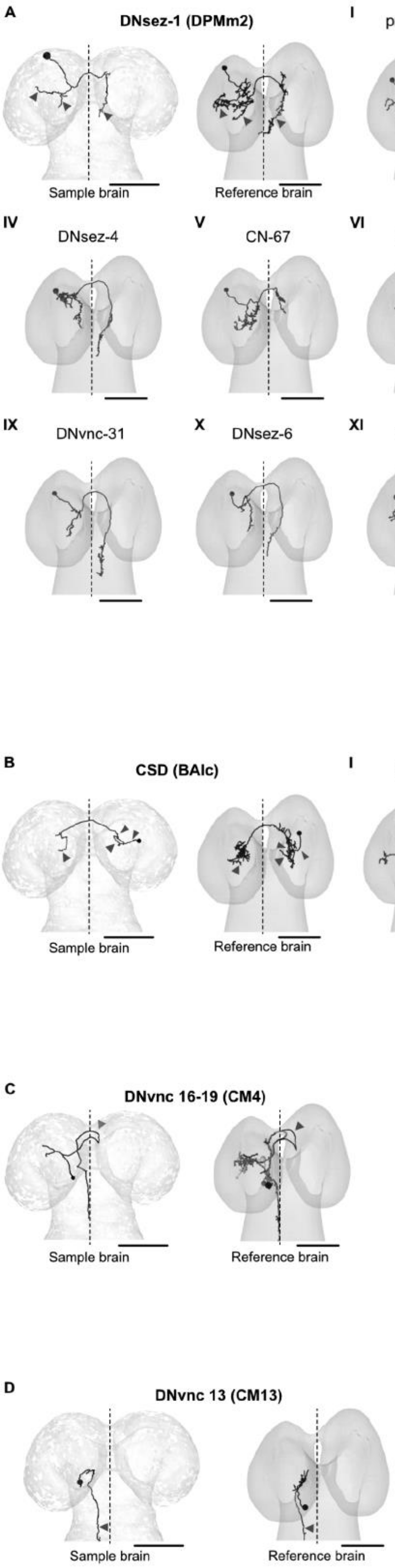

A
DNsez-1 (DPMm2)
Sample brain
Reference brain
I
pre-DNvnc-21
II
DNvnc-28
III
DNvnc-5
IV
DNsez-4
V
CN-67
VI
DNvnc-27
VII
DNsez-5
VIII
DNvnc-30
IX
DNvnc-31
X
DNsez-6
XI
DNvnc-32
B
CSD (BAlc)
Sample brain
Reference brain
I
somPN12
II
somPN11
III
somPN10
C
DNvnc 16-19 (CM4)
Sample brain
Reference brain
D
DNvnc 13 (CM13)
Sample brain
Reference brain

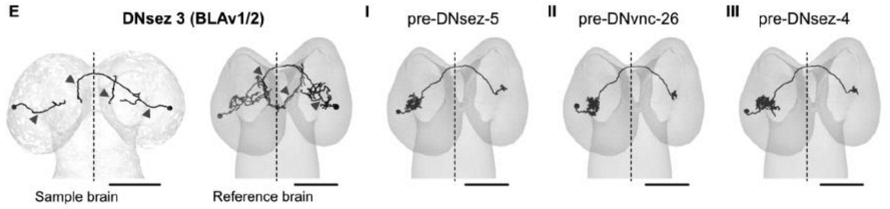
E
DNsez 3 (BLAv1/2)
I
pre-DNsez-5
II
pre-DNvnc-26
III
pre-DNsez-4
Sample brain
Reference brain

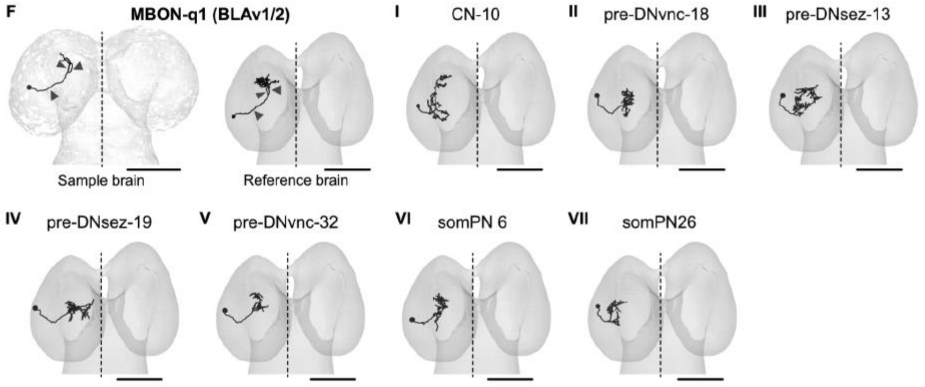
F
MBON-q1 (BLAv1/2)
I
CN-10
II
pre-DNvnc-18
III
pre-DNsez-13
Sample brain
Reference brain
IV
pre-DNsez-19
V
pre-DNvnc-32
VI
somPN 6
VII
somPN26

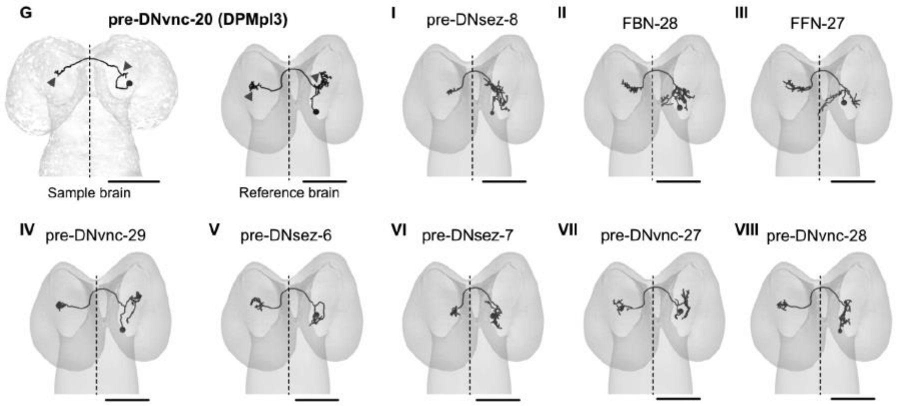
G
pre-DNvnc-20 (DPMpl3)
I
pre-DNsez-8
II
FBN-28
III
FFN-27
Sample brain
Reference brain
IV
pre-DNvnc-29
V
pre-DNsez-6
VI
pre-DNsez-7
VII
pre-DNvnc-27
VIII
pre-DNvnc-28

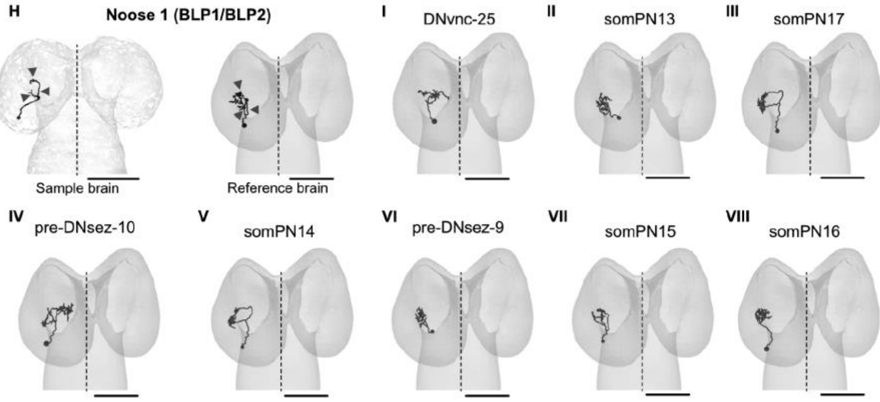
H
Noose 1 (BLP1/BLP2)
Sample brain
Reference brain
I
DNvnc-25
II
somPN13
III
somPN17
IV
pre-DNsez-10
V
somPN14
VI
pre-DNsez-9
VII
somPN15
VIII
somPN16

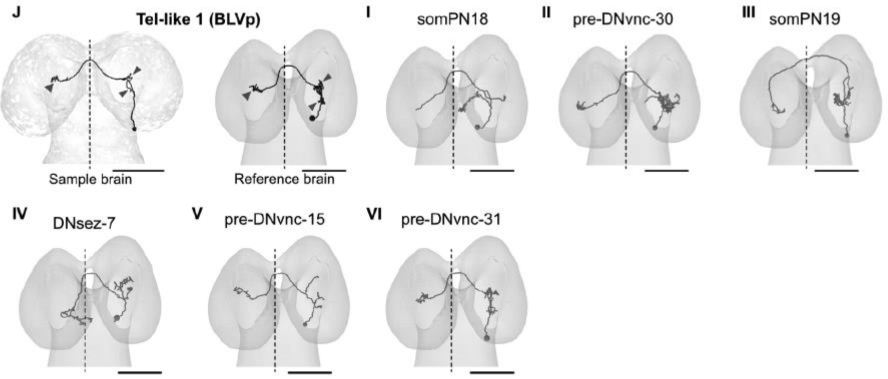
J
Tel-like 1 (BLVp)
Sample brain
Reference brain
I
somPN18
II
pre-DNvnc-30
III
somPN19
IV
DNsez-7
V
pre-DNvnc-15
VI
pre-DNvnc-31

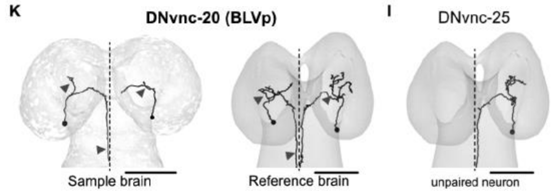
K
DNvnc-20 (BLVp)
Sample brain
Reference brain
I
DNvnc-25
unpaired neuron

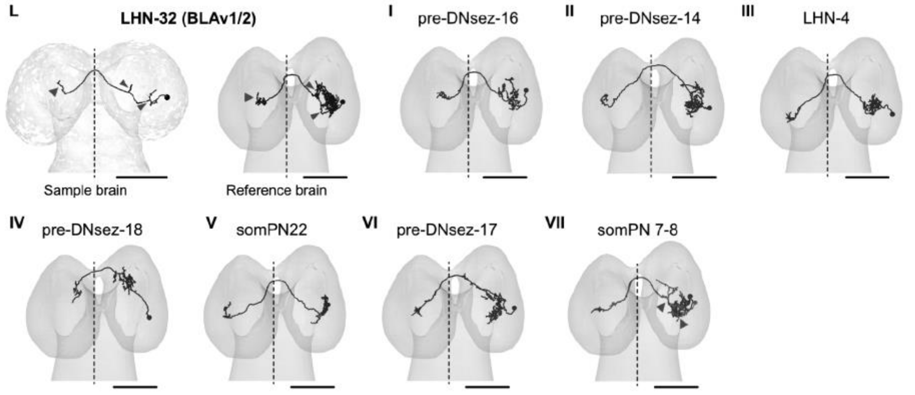
L
LHN-32 (BLAv1/2)
Sample brain
Reference brain
I pre-DNsez-16
II pre-DNsez-14
III LHN-4
IV pre-DNsez-18
V somPN22
VI pre-DNsez-17
VII somPN 7-8

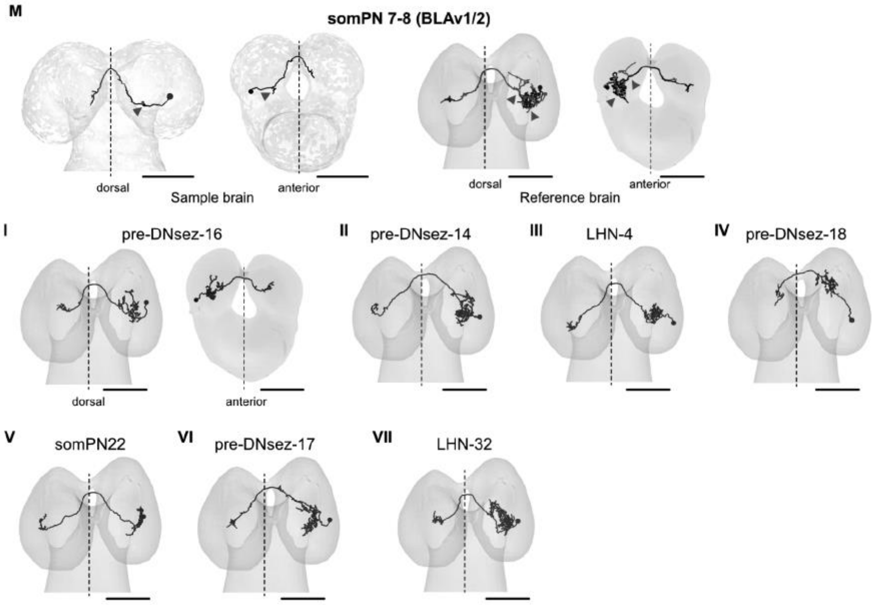
M
somPN 7-8 (BLAv1/2)
dorsal
Sample brain
anterior
dorsal
Reference brain
anterior
I pre-DNsez-16
dorsal
anterior
II pre-DNsez-14
III LHN-4
IV pre-DNsez-18
V somPN22
VI pre-DNsez-17
VII LHN-32

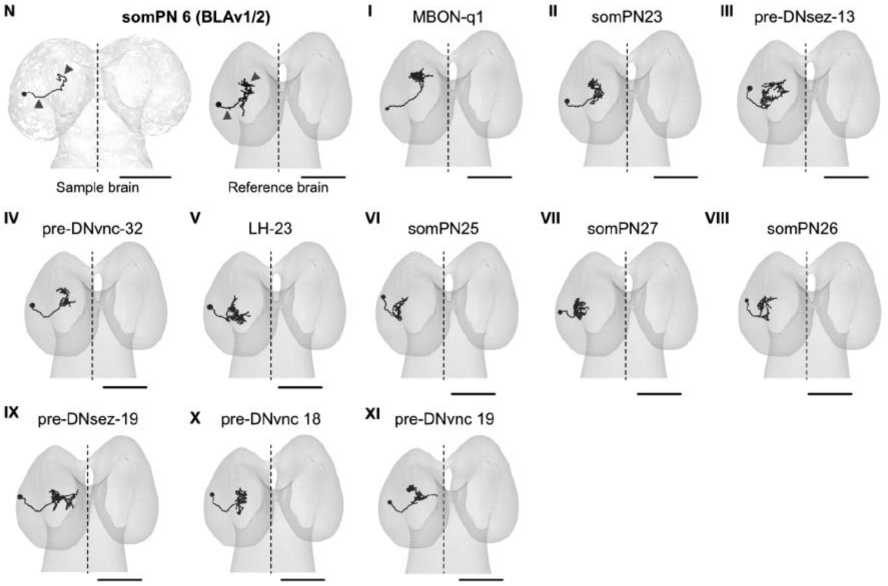
N
somPN 6 (BLAv1/2)
Sample brain
Reference brain
I MBON-q1
II somPN23
III pre-DNsez-13
IV pre-DNvnc-32
V LH-23
VI somPN25
VII somPN27
VIII somPN26
IX pre-DNsez-19
X pre-DNvnc 18
XI pre-DNvnc 19

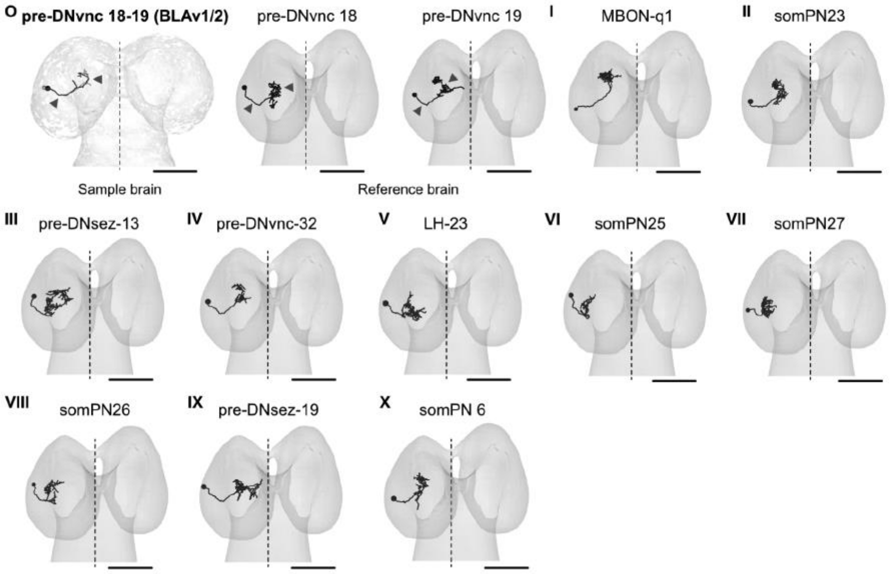
O pre-DNvnc 18-19 (BLAv1/2)
pre-DNvnc 18
pre-DNvnc 19
Sample brain
Reference brain
I MBON-q1
II somPN23
III pre-DNsez-13
IV pre-DNvnc-32
V LH-23
VI somPN25
VII somPN27
VIII somPN26
IX pre-DNsez-19
X somPN 6

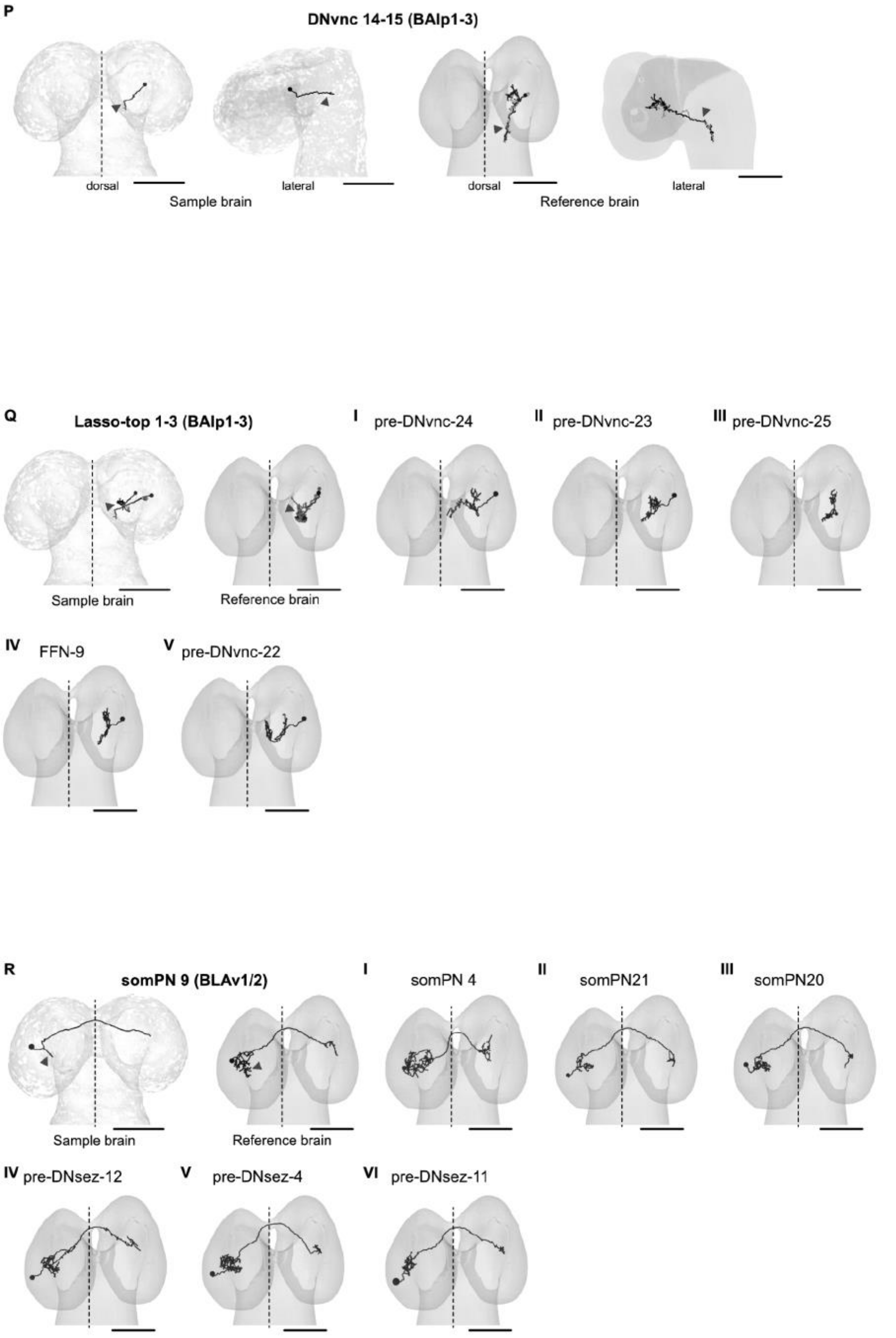

**Figure S3:** Neuron morphology atlas of identified neurons and related similar neurons from the same lineage, related to Figure 3. Arrow heads label neuron unique structures. Dashed line indicates the midline Scale: 25μm.

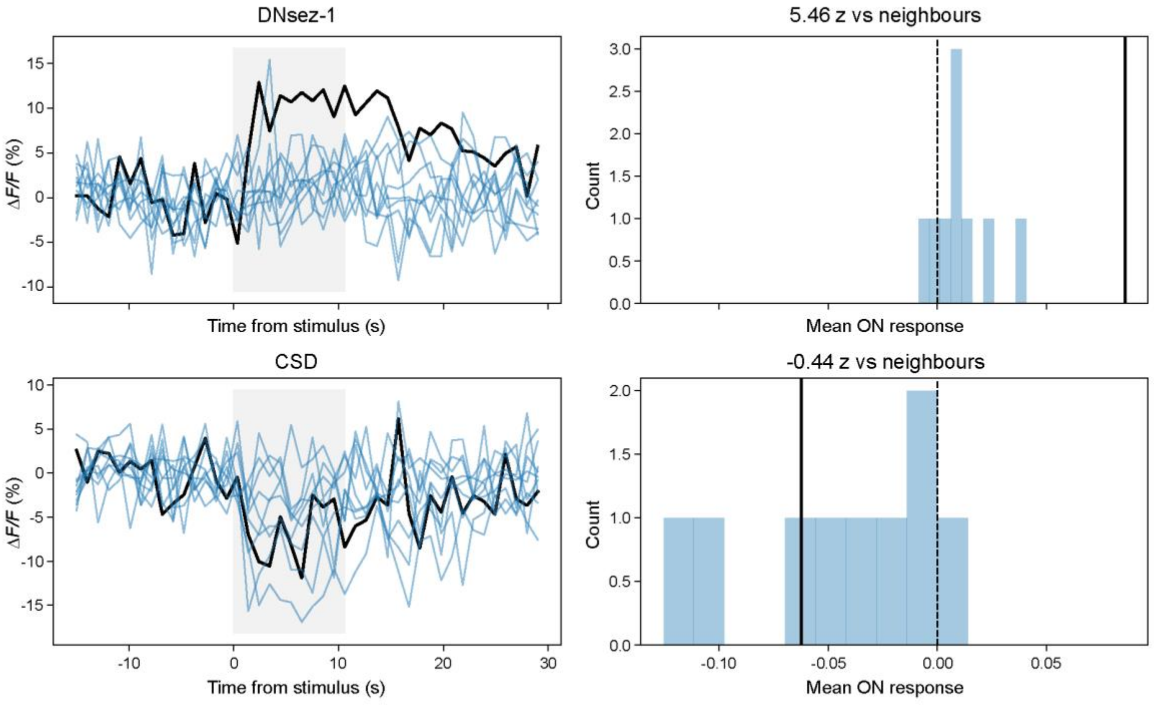

**Figure S4**. Comparison of Basin response of DNsez-1 and CSD to the responses of their nearest neighbours, related to Figure 5. Left: the response of DNsez-1 (black, top) and its ten nearest neighbours (blue, top) and CSD (black, bottom) and its ten nearest neighbours (blue, bottom) to Basin stimulation (grey shading). Right: Histogram showing comparison of the mean Basin response (DF/F) in the 10sec stimulation window of DNsez-1 (solid black line, top) or CSD (solid black line, bottom) to the distribution of responses of their ten nearest neighbours (blue). The calculated z-score for the hit neuron relative to its neighbour distribution is provided in each panel title. DNsez is a clear outlier and different to all its neighbours. CSD is amongst the 4 most inhibited neurons.

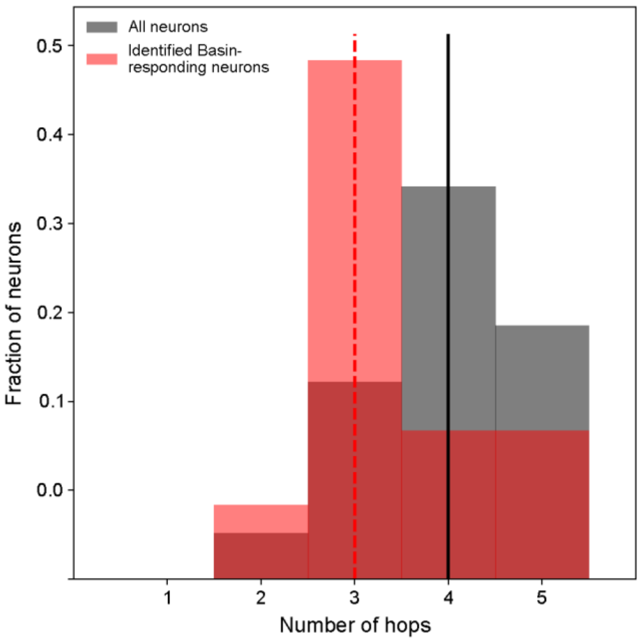

**Figure S5.** Minimum number of hops from Basin interneurons to identified Basin-responding neurons, , related to Figure 6. Normalized distribution of the shortest paths (minimum number of hops) from Basin neurons (Basins 1–4, left and right hemisegments) to identified Basin-responding neurons (red histogram) and to all brain neurons (grey histogram). The identified Basin-responding neurons have significantly shorter paths to Basins than all brain neurons (Pearson's chi-square test: $\chi^2 = 19.29$, $p < 0.0001$). Basin-responsive neurons were a median of 3 hops away from Basin neurons, while all brain neurons were a median of 4 hops away. This result indicates that Basin-responsive neurons have shorter paths to Basin neurons than would be expected by chance. Solid black and dashed red vertical lines indicate the median hop distance for all brain neurons and hit neurons, respectively.

#### Tables

**Table S1**. Cell lineages that are 2-5 hops downstream of Basins and whether they were identified in the LSM whole-brain imaging dataset, related to Figure 2.

| Cell lineage | No. of hops | Identified in functional imaging |
|---|---|---|
| BLD2-4 L | 2 | yes |
| BLAv1/2 p L | 2 | yes |
| BLVa3/4 a L | 2 | yes |
| BLVp2 L | 2 | yes |
| CP1 v L | 2 | yes |
| CP2/3 v L | 2 | yes |
| DPLp1/2 L | 2 | yes |
| DPMm2 L | 2 | yes |
| DPMpl3 L | 2 | yes |
| BLAv1/2 a L | 2 | yes |
| BLAvm p L | 2 | no |
| BLD5/6 L | 2 | no |
| BLP3/4 b L | 2 | no |
| BLVp1 L | 2 | no |
| DPMl1 L | 2 | no |
| BLAv1/2 a L | 3 | yes |
| BLAv1/2 l L | 3 | yes |
| BLAv1/2 p L | 3 | yes |
| BLD2-4 L | 3 | yes |
| BLP1/2 a L | 3 | yes |
| BLVa3/4 a L | 3 | yes |
| BLVa3/4 t L | 3 | yes |
| BLVp1 L | 3 | yes |
| BLVp2 L | 3 | yes |
| CM1/3 L | 3 | yes |
| CM4 L | 3 | yes |
| CP1 v L | 3 | yes |
| CP2/3 d L | 3 | yes |
| CP2/3 v L | 3 | yes |
| DALv2/3 L | 3 | yes |
| DPLal1-3 L | 3 | yes |
| DPLl1-3 a L | 3 | yes |
| DPMm2 L | 3 | yes |
| DPMpl3 L | 3 | yes |
| BAla1/2 L | 3 | no |
| BAla3/4 L | 3 | no |
| BAmd1 L | 3 | no |
| BAmd2 L | 3 | no |
| BAmv1/2 L | 3 | no |
| BLAd1-4 L | 3 | no |
| BLAl L | 3 | no |
| BLAvm a L | 3 | no |
| BLAvm p L | 3 | no |
| BLD5/6 L | 3 | no |
| BLP3/4 b L | 3 | no |
| BLVa1/2 a L | 3 | no |
| CP1 d L | 3 | no |
| CP4 L | 3 | no |
| DALcl1/2 v L | 3 | no |

| | | |
|---|---|---|
| DALcm1/2 m L | 3 | no |
| DALd L | 3 | no |
| DALl1 L | 3 | no |
| DALv1 L | 3 | no |
| DALv2/3 acc lat L | 3 | no |
| DALv2/3 acc med L | 3 | no |
| DALv2/3 acc ven L | 3 | no |
| DALv2/3 acc ven lat L | 3 | no |
| DPLam L | 3 | no |
| DPLc4 L | 3 | no |
| DPLc5 a L | 3 | no |
| DPLd L | 3 | no |
| DPMl1 L | 3 | no |
| DPMl2 L | 3 | no |
| DPMl3/4 p L | 3 | no |
| DPMm1 L | 3 | no |
| DPMpl1/2 L | 3 | no |
| BAlc L | 4 | yes |
| BAlp1-3 L | 4 | yes |
| BLAv1/2 a L | 4 | yes |
| BLAv1/2 l L | 4 | yes |
| BLAv1/2 p R | 4 | yes |
| BLD2-4 L | 4 | yes |
| BLP1/2 a L | 4 | yes |
| BLP1/2 b L | 4 | yes |
| BLVa3/4 a L | 4 | yes |
| BLVa3/4 b L | 4 | yes |
| BLVa3/4 t L | 4 | yes |
| BLVp1 L | 4 | yes |
| BLVp2 L | 4 | yes |
| CM1/3 L | 4 | yes |
| CM4 L | 4 | yes |
| CP1 v L | 4 | yes |
| CP2/3 d L | 4 | yes |
| CP2/3 v L | 4 | yes |
| DALv2/3 L | 4 | yes |
| DPLal1-3 L | 4 | yes |
| DPLl1-3 a L | 4 | yes |
| DPLp1/2 L | 4 | yes |
| DPMm2 L | 4 | yes |
| DPMpl3 L | 4 | yes |
| BAla1/2 L | 4 | no |
| BAla3/4 L | 4 | no |
| BAmd1 L | 4 | no |
| BAmd2 L | 4 | no |
| BAmv1/2 L | 4 | no |
| BLAd1-4 L | 4 | no |
| BLAl L | 4 | no |
| BLAv1/2 v L | 4 | no |
| BLAvm a L | 4 | no |
| BLAvm c L | 4 | no |
| BLAvm p L | 4 | no |
| BLD1 L | 4 | no |
| BLP3/4 a L | 4 | no |
| BLP3/4 b L | 4 | no |
| BLVa1/2 a L | 4 | no |
| BLVa1/2 b L | 4 | no |

| | | |
|---|---|---|
| CP1 d L | 4 | no |
| CP4 L | 4 | no |
| DALcl1/2 d L | 4 | no |
| DALcl1/2 v L | 4 | no |
| DALcm1/2 m L | 4 | no |
| DALcm1/2 v L | 4 | no |
| DALd L | 4 | no |
| DALl1 L | 4 | no |
| DALv1 L | 4 | no |
| DALv2/3 acc lat L | 4 | no |
| DALv2/3 acc med L | 4 | no |
| DALv2/3 acc ven L | 4 | no |
| DALv2/3 acc ven lat L | 4 | no |
| DAMd1 L | 4 | no |
| DAMd2/3 L | 4 | no |
| DAMv1/2 L | 4 | no |
| DILP L | 4 | no |
| DPLam L | 4 | no |
| DPLc1 L | 4 | no |
| DPLc3 L | 4 | no |
| DPLc4 L | 4 | no |
| DPLc5 a L | 4 | no |
| DPLc5 p L | 4 | no |
| DPLd L | 4 | no |
| DPLl1-3 i L | 4 | no |
| DPLl1-3 L | 4 | no |
| DPLm1 L | 4 | no |
| DPLm2 L | 4 | no |
| DPMl1 L | 4 | no |
| DPMl2 L | 4 | no |
| DPMl3/4 p L | 4 | no |
| DPMm1 L | 4 | no |
| DPMpl1/2 L | 4 | no |
| DPMpm1 L | 4 | no |
| DPMpm2 L | 4 | no |
| BAlc L | 5 | yes |
| BAlp1-3 L | 5 | yes |
| BAmas1/2 L | 5 | yes |
| BLAv1/2 a L | 5 | yes |
| BLAv1/2 l R | 5 | yes |
| BLAv1/2 p L | 5 | yes |
| BLD2-4 L | 5 | yes |
| BLP1/2 a L | 5 | yes |
| BLP1/2 b L | 5 | yes |
| BLVa3/4 a L | 5 | yes |
| BLVa3/4 b L | 5 | yes |
| BLVa3/4 t L | 5 | yes |
| BLVp1 L | 5 | yes |
| BLVp2 L | 5 | yes |
| CM1/3 L | 5 | yes |
| CM4 L | 5 | yes |
| CP1 v L | 5 | yes |
| CP2/3 d L | 5 | yes |
| CP2/3 v L | 5 | yes |
| DALv2/3 L | 5 | yes |
| DPLal1-3 L | 5 | yes |
| DPLl1-3 a L | 5 | yes |

| | | |
|---|---|---|
| DPLp1/2 L | 5 | yes |
| DPMm2 L | 5 | yes |
| DPMpl3 L | 5 | yes |
| KeC L | 5 | yes |
| BAla1/2 L | 5 | no |
| BAla3/4 L | 5 | no |
| BAlp4 L | 5 | no |
| BAlv L | 5 | no |
| BAmd1 L | 5 | no |
| BAmd2 L | 5 | no |
| BAmv1/2 L | 5 | no |
| BAmv3 L | 5 | no |
| BLAd1-4 L | 5 | no |
| BLAl L | 5 | no |
| BLAvm a L | 5 | no |
| BLAvm c L | 5 | no |
| BLD1 R | 5 | no |
| BLP3/4 a L | 5 | no |
| BLP3/4 b L | 5 | no |
| BLVa1/2 a L | 5 | no |
| BLVa1/2 b L | 5 | no |
| CP1 d L | 5 | no |
| DALcl1/2 v L | 5 | no |
| DALcm1/2 v L | 5 | no |
| DALd L | 5 | no |
| DALl1 L | 5 | no |
| DALv1 L | 5 | no |
| DALv2/3 acc ven L | 5 | no |
| DAMd1 L | 5 | no |
| DAMv1/2 L | 5 | no |
| DILP L | 5 | no |
| DPLam L | 5 | no |
| DPLc1 L | 5 | no |
| DPLc4 L | 5 | no |
| DPLc5 p L | 5 | no |
| DPLd L | 5 | no |
| DPLl1-3 i L | 5 | no |
| DPLm1 L | 5 | no |
| DPLm2 L | 5 | no |
| DPMl1 L | 5 | no |
| DPMl1 R | 5 | no |
| DPMl2 L | 5 | no |
| DPMl3/4 a L | 5 | no |
| DPMl3/4 p L | 5 | no |
| DPMm1 L | 5 | no |
| DPMpl1/2 L | 5 | no |
| DPMpm1 L | 5 | no |
| DPMpm2 L | 5 | no |

**Table S2.** Identified brain neurons with their corresponding cell lineage, cell type, sensory modality, hub-neuron identity, and synaptic connectivity, related to Figure 3 (synaptic threshold: 0.01; *synaptic threshold: ≥3 synapses). Candidate neurons, which could not be uniquely identified, are listed individually. Hub neurons are defined as having ≥20 in- or out-degree[1].

| skid Reference brain | skid Sample brain | Publication name | Cell lineage | Cell type | Sensory modality | Synaptic connectivity downstream Basin | hub neuron |
|---|---|---|---|---|---|---|---|
| 3044500 | 26235 | DNsez-1_stop | DPMm2 | DN-SEZ | 3rd order: noci, mechano | 3-hop | yes |
| 9234420 | 23023 | DNsez-3 | BLAv1/2 | DN-SEZ | 3rd order: noci, mechano | 3-hop | yes |
| 17728922 | 47849 | DNsez-3 | BLAv1/2 | DN-SEZ | 3rd order: noci, mechano | 3-hop | yes |
| 8723983 | 49438 | DNvnc-13 | CM1/3 | DN-VNC | 4th order: gustatory, olfactory | 5-hop | no |
| 17075832 | 75841 | DNvnc-14 | BAlp 1-3 | DN-VNC | 3rd order: gustatory | 4-hop | no |
| 7439913 | 75841 | DNvnc-15 | BAlp 1-3 | DN-VNC | 3rd order: gustatory | 4-hop | no |
| 10915553 | 54086 | DNvnc-16 | CM4 | DN-VNC | 3rd order: noci | 3-hop | no |
| 10609443 | 54086 | DNvnc-17 | CM4 | DN-VNC | 3rd order: noci, mechano | 3-hop | no |
| 17732270 | 54086 | DNvnc-18 | CM4 | DN-VNC | 3rd order: mechano | 4-hop | no |
| 7227010 | 54086 | DNvnc-19 | CM4 | DN-VNC | 3rd order: noci | 3-hop | no |
| 10945795 | 373620 | DNvnc-20 | BLVp2 | DN-VNC | 3rd order: noci, mechano | 3-hop | no |
| 15588711 | 30208 | DNvnc-20 | BLVp2 | DN-VNC | 3rd order: noci, mechano | 3-hop | no |
| N/A | 32958 | Kenyon cells | KC | KC | N/A | 5-hop* | no |
| N/A | 32966 | Kenyon cells | KC | KC | N/A | 5-hop* | no |
| N/A | 82755 | Kenyon cells | KC | KC | N/A | 5-hop* | no |
| N/A | 403750 | Kenyon cells | KC | KC | N/A | 5-hop* | no |
| N/A | 413009 | Kenyon cells | KC | KC | N/A | 5-hop* | no |
| 5118060 | 23027 | LHN-32 | BLAv1/2 | LHNs | 2nd mechano, 3rd olfactory | 4-hop | yes |
| 7910624 | 29643 | MBON-q1 | BLAv1/2 | MBON | 3rd order: mechano, respiratory | 5-hop | no |
| 17728723 | 395228, 395507 | Lasso-top 1 | BAlp 1-3 | N/A | 3rd order: gustatory | 7-hop | no |
| 17728730 | 395228, 395507 | Lasso-top 2 | BAlp 1-3 | N/A | 3rd order: gustatory | 5-hop | no |
| 8809171 | 395228, 395507 | Lasso-top 3 | BAlp 1-3 | N/A | 4th order: noci, gustatory, olfactory, thermio-warm | 5-hop | no |
| 14270761 | 58156 | Tel-like 1 | BLVp1 | N/A | 3rd order: noci | 3-hop | no |
| 5744502 | 405426 | Noose 1 | BLVp2 | N/A | 3rd order: mechano | 4-hop | no |
| 16541149 | 29651 | somPN 6 | BLAv1/2 | PN-somato | 2nd order: noci | 2-hop | no |
| 6623089 | 22908, 8925 | pre-DNvnc-18 | BLAv1/2 | pre-DN-VNC | 2nd order: noci | 3-hop | no |
| 16931001 | 22908, 8925 | pre-DNvnc-19 | BLAv1/2 | pre-DN-VNC | 3rd order: noci, mechano | 3-hop | no |
| 11291653 | 395927 | somPN 7 | BLAv1/2 | PN-somato | 2nd order: noci | 4-hop | no |
| 11263687 | 395927 | somPN 8 | BLAv1/2 | PN-somato | 2nd order: noci | 2-hop | yes |
| 20730780 | 22916 | somPN 9 | BLAv1/2 | PN-somato | 2nd order: noci | 2-hop | no |
| 4966994 | 32853 | pre-DNvnc-20 | DPMpl3 | pre-DN-VNC | 3rd order: noci, propio | 3-hop | yes |
| 13849557 | 395150 | CSD | BALc | pre-DN-VNC | 3rd order: gustatory, mechano, olfactory, thermo-warm | 4-hop | yes |

**Table S3.** Skeleton IDs (skids) and names of previously published neurons.

| Left skid | Right skid | Neuron name |
|---|---|---|
| 1678567 | 6153877 | DNvnc-21 |
| 10728328 | 18464581 | DNvnc-22 |
| 17165755 | 13727202 | DNvnc-23 |
| 16851496 | 16339338 | DNvnc-24 |
| 5763407 | 10945111 | DNvnc-25 |
| 15365184 | 5512988 | DNvnc-26 |
| 13749806 | 14034206 | DNvnc-27 |
| 8834812 | 18970354 | A02 |
| 7766185 | 7934152 | SeIN134 |
| 1767828 | 4139768 | A12m |
| 4081479 | 5336476 | A19c |
| 4740178 | 4833168 | A05q |
| 21250110 | 9640873 | A10j |
| 4680995 | 17806341 | LHN-4 |
| 14319185 | 8565131 | CN-57 |
| 16486385 | 11178891 | CN-46 |
| 10540188 | 8932110 | Cheech |
| 12498338 | 3913629 | CN-50 |
| 12476851 | 16070049 | CN-1 |
| 17434825 | 16485079 | CN-47 |
| 3044500 | 6317793 | DNsez-1_stop |
| 17728922 | 9234420 | DNsez-3 |
| 8723983 | 16115255 | DNvnc-13 |
| 17075832 | 14039007 | DNvnc-14 |
| 7439913 | 15934458 | DNvnc-15 |
| 10915553 | 11905911 | DNvnc-16 |
| 10609443 | 6114695 | DNvnc-17 |
| 17732270 | 6125806 | DNvnc-18 |
| 7227010 | 16115575 | DNvnc-19 |
| 15588711 | 10945795 | DNvnc-20 |
| 5118060 | 15605987 | LHN-32 |
| 7910624 | 7897469 | MBON-q1 |
| 17728723 | 13084891 | Lasso-top 1 |
| 17728730 | 17629213 | Lasso-top 2 |
| 8809171 | 18187865 | Lasso-top 3 |
| 14270761 | 9103710 | Tel-like 1 |
| 5744502 | 1805406 | Noose 1 |
| 16541149 | 10079731 | somPN 6 |
| 6623089 | 17897230 | pre-DNvnc-18 |
| 16931001 | 8709865 | pre-DNvnc-19 |
| 11291653 | 8412625 | somPN 7 |
| 11263687 | 6692908 | somPN 8 |
| 20730780 | 9202326 | somPN 9 |
| 4966994 | 16578252 | pre-DNvnc-20 |
| 13849557 | 6143930 | CSD |
| 3043970 | 16181130 | pre-DNvnc-21 |
| 3048606 | 3009700 | DNvnc-28 |

| | | |
|---|---|---|
| 3946166 | 3620633 | DNvnc-5 |
| 9083312 | 14077179 | DNsez-4 |
| 11216417 | 8451818 | CN-67 |
| 16340977 | 18232369 | DNsez-5 |
| 16561660 | 13652117 | DNvnc-30 |
| 17102002 | 1065967 | DNvnc-31 |
| 17321919 | 4215611 | DNsez-6 |
| 17473163 | 16571210 | DNvnc-32 |
| 11934107 | 3844961 | somPN10 |
| 11986594 | 6218461 | somPN11 |
| 12121795 | 11788830 | somPN12 |
| 8802069 | 4665627 | FFN-9 |
| 17383431 | 9532295 | pre-DNvnc-22 |
| 17728695 | 10367190 | pre-DNvnc-23 |
| 12425856 | 13173311 | pre-DNvnc-24 |
| 18106257 | 17031293 | pre-DNvnc-25 |
| 16472086 | 16433177 | pre-DNsez-4 |
| 16594752 | 530532 | pre-DNvnc-26 |
| 21220994 | 8229118 | pre-DNsez-5 |
| 14493841 | 11361875 | FFN-27 |
| 18538744 | 11362334 | pre-DNvnc-27 |
| 15028907 | 11362589 | pre-DNsez-6 |
| 10612094 | 12122924 | FBN-28 |
| 10248031 | 17380319 | pre-DNvnc-28 |
| 14999928 | 4591713 | pre-DNvnc-29 |
| 5902352 | 5936935 | pre-DNsez-7 |
| 18023286 | 9753494 | pre-DNsez-8 |
| 15586949 | 3180681 | pre-DNsez-9 |
| 17105977 | 8970595 | somPN13 |
| 3819859 | 4127944 | somPN14 |
| 5747036 | 20443544 | pre-DNsez-10 |
| 5767836 | 20443505 | somPN15 |
| 17468727 | 127938 | somPN16 |
| 19929273 | 20479959 | somPN17 |
| 19101351 | 3848843 | somPN18 |
| 17729088 | 9637388 | pre-DNvnc-30 |
| 19884188 | 10945581 | somPN19 |
| 5764924 | 10947427 | DNsez-7 |
| 17804729 | 11050498 | pre-DNvnc-15 |
| 18035063 | 17937986 | pre-DNvnc-31 |
| 4663875 | 11271082 | pre-DNsez-11 |
| 9508724 | 17541953 | somPN20 |
| 10282065 | 5946241 | pre-DNsez-12 |
| 19102929 | unpaired | somPN21 |
| 21136945 | 17654638 | somPN 4 |
| 21220994 | 8229118 | pre-DNsez-14 |
| 5657004 | 5134110 | pre-DNsez-16 |
| unpaired | 17588623 | somPN22 |

| | | |
|---|---|---|
| 10282065 | 5946241 | pre-DNsez-17 |
| 12364717 | 11356523 | pre-DNsez-18 |
| 17108278 | unpaired | somPN23 |
| 13714311 | 8710444 | pre-DNvnc-32 |
| 16100461 | 8711574 | LH-23 |
| 16141217 | 13722516 | somPN25 |
| 17513284 | 17361566 | somPN26 |
| 18111410 | 4876839 | pre-DNsez-19 |
| 20981350 | 8715940 | somPN27 |
| 10893103 | 9720614 | pre-DNsez-13 |
| 17150791 | 8712361 | CN-10 |
| 15547235 | N/A | HT3A |
| 15547235 | 17036160 | PN2 |
| 10160250 | 3450751 | mPN 5 |
| 3034133 | 3282869 | Basin-1 |
| 3041612 | 3074106 | Basin-2 |
| 10179501 | 3091943 | Basin-3 |
| 3040481 | 4049878 | Basin-4 |

**Table S4.** Identified neurons per lineage pair.

| Lineage | No. of Neurons (Sample Brain) | No. of Neurons (Reference Brain) | % |
|---|---|---|---|
| BLP1/2 | 4 | 102 | 3.9 |
| BLVa3/4 | 9 | 99 | 9.1 |
| BLAv1/2 | 30 | 111 | 27.0 |
| CM1/3 | 1 | 95 | 1.1 |
| CM4 | 4 | 184 | 2.2 |
| CP1 v | 1 | 62 | 1.6 |
| CP2/3v | 1 | 80 | 1.3 |
| DALv 2/3 | 1 | 42 | 2.4 |
| DPLal 1-3 | 2 | 82 | 2.4 |
| DPLl 1-3a | 1 | 24 | 4.2 |
| DPLp 1/2 | 4 | 26 | 15.4 |
| DPMm2 | 2 | 46 | 4.3 |
| CP2/3 d | 1 | 48 | 2.1 |
| DPMpl3 | 1 | 30 | 3.3 |
| KC | 5 | 223 | 2.2 |
| BLVp1, BLVp2 | 12 | 62 | 19.4 |
| Balc | 1 | 86 | 1.2 |
| BLD2-4 | 17 | 72 | 23.6 |
| BAlp1-3 | 3 | 68 | 4.4 |
| BAmas1/2 | 1 | 98 | 1.0 |
| **Sum** | 101 | 1640 | |
| **Mean** | 5.05 | 82 | 6.6 |
| **Median** | 2 | 76 | 2.9 |

**Table S5.** Number of neurons 2-hop downstream of basins per lineage pair.

| Lineage | No. of Neurons | 2-hop downstream of basins | % |
|---|---|---|---|
| BLAv1/2 | 111 | 29 | 26.1 |
| BLVa3/4 | 99 | 20 | 20.2 |
| BLVp1, BLVp2 | 62 | 14 | 22.6 |
| BLD2-4 | 72 | 8 | 11.1 |
| BLP1/2 | 102 | 6 | 5.9 |
| CM1/3 | 95 | 2 | 2.1 |
| CM4 | 184 | 3 | 1.6 |
| CP1 v | 62 | 10 | 16.1 |
| CP2/3v | 80 | 4 | 5.0 |
| DALv 2/3 | 42 | 0 | 0.0 |
| DPLal 1-3 | 82 | 13 | 15.9 |
| DPLl 1-3a | 24 | 0 | 0.0 |
| DPLp 1/2 | 26 | 3 | 11.5 |
| DPMm2 | 46 | 4 | 8.7 |
| CP2/3d | 48 | 0 | 0.0 |
| DPMpl3 | 30 | 3 | 10.0 |
| KC | 223 | 0 | 0.0 |
| Balc | 86 | 1 | 1.2 |
| BAlp1-3 | 68 | 0 | 0.0 |
| BAmas1/2 | 98 | 3 | 3.1 |
